# Modelling homing suppression gene drive in haplodiploid organisms

**DOI:** 10.1101/2021.10.12.464047

**Authors:** Yiran Liu, Jackson Champer

## Abstract

Gene drives have shown great promise for suppression of pest populations. These engineered alleles can function by a variety of mechanisms, but the most common is the CRISPR homing drive, which converts wild-type alleles to drive alleles in the germline of heterozygotes. Some potential target species are haplodiploid, in which males develop from unfertilized eggs and thus have only one copy of each chromosome. This prevents drive conversion, a substantial disadvantage compared to diploids where drive conversion can take place in both sexes. Here, we study homing suppression gene drives in haplodiploids and find that a drive targeting a female fertility gene could still be successful. However, such drives are less powerful than in diploids and suffer more from functional resistance alleles. They are substantially more vulnerable to high resistance allele formation in the embryo due to maternally deposited Cas9 and gRNA and also to somatic cleavage activity. Examining spatial models where organisms move over a continuous landscape, we find that haplodiploid suppression drives surprisingly perform nearly as well as in diploids, possibly due to their ability to spread further before inducing strong suppression. Together, these results indicate that gene drive can potentially be used to effectively suppress haplodiploid populations.

## Introduction

Suppression gene drives have recently shown great promise for a variety of applications, in particular elimination of pest populations^1–4^. These genetic elements bias their own inheritance to increase in frequency in a population, eventually causing suppression by biasing the sex ratio or rendering enough individuals sterile or nonviable. Though the most rapid progress has been in *Anopheles* mosquitoes for reduction of vector-borne disease^5–7^ (for which modification drives have also been developed^8–10^), suppression gene drives could also be used to remove invasive species or agricultural pests. Thus far, gene drives have been demonstrated in a variety of organisms, including yeast^11,12^, flies^13–15,16–18^, mice^19^, and plants^20^. Most of these have been homing types drives, where a nuclease, usually CRISPR/Cas9, cuts a wild-type chromosome at a site directed by its guide RNA (gRNA)^21^. The chromosome then undergoes homology-directed repair, which results in the drive alleles being copied to the target site. Since this occurs in the germline, the drive alleles will be inherited at an increased rate. However, if end-joining repair occurs instead, than the wild-type site can mutate into a resistance allele, which cannot be converted to a drive allele^22–24^. These resistance alleles have the potential to slow or even stop the spread of a gene drive. The dynamics of such drives have been modelled extensively^16,25–29^, which is particularly important for predicting the outcome of a real-world drive release.

Though many possible applications of gene drives have been considered^1–4^, haplodiploid species have received little attention as possible targets. In these species, males develop from unfertilized eggs and thus only have one of each chromosome, which limits drive conversion in homing drives to females. So far, there has only been a single modelling studying assessing the possibilities for modification drives in haplodiploids^30^. The study concluded that homing drives could be effective, but they were always slower than in diploids, especially when drive performance was reduced. Though these results are promising in some situations, rapid suppression of haplodiploids is often desirable. This is especially true in well-known pest species such as invasive fire ants, which can cause human harm in addition to catastrophic damage to ecosystems and agricultural production^31,32^. In this species, CRISPR genome engineering has already been demonstrated^33^, bringing the possibility of developing a gene drive closer to reality. Other major haplodiploid pests that could potentially be the target of a suppression drive include certain bark beetles, which are highly invasive and damaging to many natural forests and the lumber industry, and thrips, which cause major agricultural losses of potatoes, onions, cotton, and other crops.

However, several drive mechanisms in diploids would not function in haplodiploids, particularly those that bias sex ratio or cause infertility or nonviability in males or in both males and females. Two studies attempted to develop models for haplodiploid suppression drives based on homing drives, but these lacked power to cause complete suppression on their own^34,35^. Our initial assessments indicated that of the established drive systems, only homing drives targeting haplosufficient but essential female fertility genes could cause strong suppression. In this study, we investigate this drive in detail, including its properties such as rate of spread and especially its overall suppressive power, comparing it to more well-studied but similar suppression drives in diploids. We find that embryo resistance and somatic cleavage activity can substantially reduce the efficiency of haplodiploid suppression drives, but over a wide parameter range, its performance is still nearly as good as diploid drives.

Additionally, many recent modelling studies of gene drive suppression have revealed unexpected complexity in spatially explicit scenarios. These scenarios could take the form of abstract patches^36^, complex networks of linked panmictic populations^37,38^, or arenas with continuous space^39–41^. In these simulations, the gene drive is often initially successful, eliminating the population in a particular region. However, this region can then be recolonized by wild-type individuals. The gene drive is still present in adjacent areas and moves in shortly afterward, completing cycles of suppression and recolonization as the drive “chases” wild-type alleles. Because even slightly weaker drives are more prone to suffering from such outcomes^39^, a haplodiploid homing suppression drive might lose its ability to effectively suppress populations. We used our continuous space model to assess the performance of haplodiploid drives, but we found that it was only, on average, slightly more vulnerable to chasing than diploid drives of the same design and performance, still usually suppressing the population in a timely manner. In some areas of parameter space, the haplodiploid drive even had small advantages over the diploid version. These results indicate that homing suppression gene drives targeting female fertility genes are promising candidates for genetic control of haplodiploid pest species.

## Methods

### Suppression drive strategy

In haplodiploids, females are derived from the diploid progeny, and unfertilized eggs become males. Here, we assess homing drives targeting female fertility genes for population suppression. In drive heterozygous females (males with only one chromosome cannot be heterozygous), the wild-type alleles will be converted into drive alleles in the germline through target cleavage followed by homology-directed repair using the drive as a template. This is the “drive conversion rate” or “drive efficiency”. With such a drive, we target a haplosufficient but essential female fertility gene, leading to sterility in female drive homozygotes. The increasing frequency of drive alleles will result in accumulation of sterile drive homozygous females among the population, which can ultimately cause the population collapse.

However, there are two possible repair mechanisms after DNA cleavage, homology-directed repair and end-joining. If the cleaved target site is repaired by the latter mechanism, it may cause the site to be mutated, and such sites couldn’t be cleaved by Cas9 in the future due to sequence mismatch with the gRNA. Because these can no longer be converted to drive alleles, they are called “resistance alleles”. Most resistance alleles result in a non-functional target gene, which we called “r2” alleles. Such alleles could contribute to female sterility but would also slow the drive. Females possessing only drive and/or nonfunctional resistance alleles are considered to be sterile in our model. Functional “r1” resistance alleles, a more serious type, will almost always result in failure of a suppression drive. However, multiple gRNAs^16,42^ and conserved target sites^5^ could be used to limit the formation of functional resistance alleles (perhaps at the cost of drive conversion efficiency^16,42,43^), so we only explicitly modelled them in some of our simulations.

### Simulation model

The forward-in-time genetic simulation framework SLiM (version 3.6) was used to perform all simulations^44^. In this study, we used code for diploids targeting X-linked loci to model haplodiploids since inheritance patterns in both situations are identical. The model is described in detail in a previous study^39^. In short, the sexually reproducing population in our model is 50,000 with a 1:1 sex ratio and discrete generations. This means that for each time step in the model, completely new individuals are generated in a single round of reproduction using density-dependent fecundity, with no individuals persisting from one generation to the next. Drive conversion and various fitness effects are modelled explicitly. In the spatial model, individual migrate in even-field, bounded, dimensionless continuous space, experience density effects, finding mates, and distributing offspring in only their local area. See the Supplemental Methods for full details of the models.

## Results

### Characteristics of haplodiploid suppression drives

To assess the potential of a female fertility homing suppression drive in haplodiploids, we first compared drive characteristics between haplodiploids and diploids in panmictic populations. Our model uses discrete generations and simplified competition and lifecycle characteristics. Competition is simply determined by the size of the total population compared to the carrying capacity, which affects female fecundity. Thus, rather than being representative of a particular species, it provides an easy way to focus on comparisons between drives with different performance parameters while still retaining minimal ecological features to ensure that the population has a stable carrying capacity.

Our default parameters (Table S1) approximately corresponded to a successfully constructed drive in *Anopheles* with reduced fitness costs^5^, representing highly efficient but somewhat imperfect systems. The suppression drive targets a female fertility gene, causing females without wild-type alleles to be sterile (Figure 1A). In diploids, wild-type alleles could be converted into drive alleles during the germline stage in both males and females (Figure 1A), so that the drive frequency could increase rapidly (Figure 1B). Males have only one chromosome in haplodiploids, so drive conversion can only be completed in females (Figure 1A), resulting in a slower rate of increase (Figure 1B). Thus, the haplodiploid drive could still eliminate the population, but this occurred later than in diploids (Figure 1C). This delay was exacerbated in a small number of simulations in which stochastic effects allowed the wild-type population to temporarily rebound before ultimately being suppressed.

**Figure 1.**
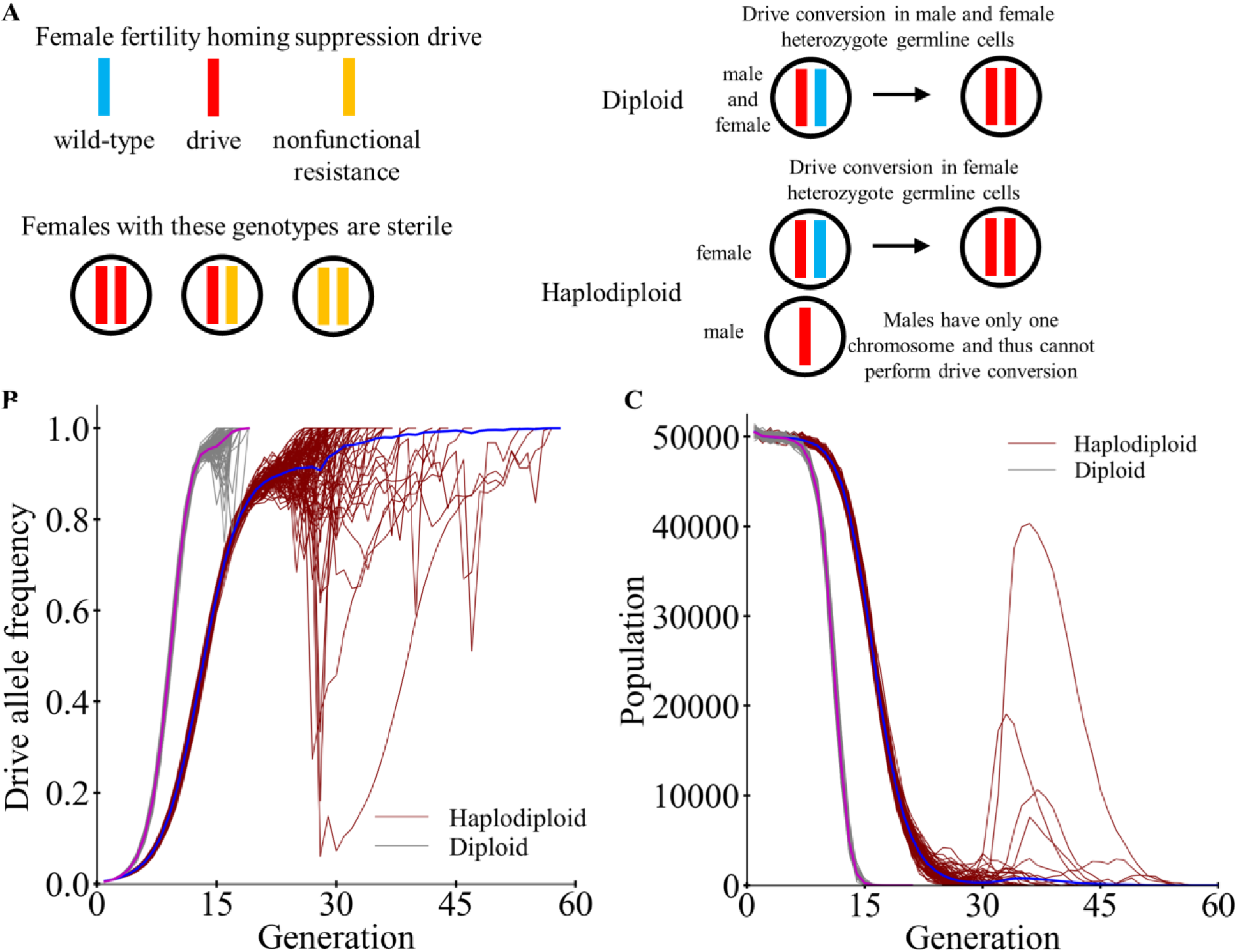
Homing suppression drives in haplodiploid and diploid populations. (A) In a female fertility homing drive, females with only drive alleles or nonfunctional resistance alleles are sterile. Such nonfunctional resistance alleles can be formed in the germline as an alternative to drive conversion or in the embryo from matern ally deposited Cas9 and gRNA. The drive can also be impeded by somatic cleavage activity, which can reduce fertility in drive/wild-type heterozygotes. In diploids, the drive can increase in frequency quickly since drive conversion can take place in the germline of male and female heterozygotes. In haplodiploids, males form from unfertilized eggs and thus only have a single chromosome, allowing drive conversion only in females. (**B,C**) Drive heterozygotes (and drive males for the haplodiploid drive) with default performance characteristics were released at 1% frequency into a panmictic population of wild-type individuals. The (**B**) drive allele frequency and (**C**) population size were tracked for each generation. Each thin line shows the frequency and population trajectories for an individual simulation (100 total for each drive), and thick lines show the average values.

Aside from affecting the rate of increase of the drive (Figure S1 and accompanying text), various performance parameters can also affect genetic load, which is a characteristic that measures suppressive power on a target population. It measures the reduction in reproductive capacity in the population compared to an equivalent wild-type population. A genetic load of 0 means that the drive does not affect reproduction, while a genetic load of 1 means that the drive has stopped all reproduction in the population. Higher genetic loads will generally reduce the population size to a greater degree, but the exact level of reduction depends on complex species-specific and ecology-specific density dependence and other related factors^3^. Thus, when assessing the power of a suppression drive, genetic load provides a convenient measure to examine effects of the drive itself. Note that a drive with sufficiently high genetic load will have enough power to eliminate the population if the low-density growth rate is sufficiently low, unless other factors intervene such as functional resistance alleles or spatial structure.

We measured the genetic load of the suppression drives after a large between generations 50-150, at which point they had reached their long-term equilibrium frequency (Figure S2A). In general, the genetic load of diploid drives was higher than for haplodiploids (Figure 2), though the haplodiploid drive was still able to reach high genetic load at high efficiency (Figure 2A, S2B). Variation in drive efficiency (Figure 2A) and fitness (Figure 2B) had similar effects on both drives, though the effect was somewhat larger in haplodiploids. However, the genetic load in haplodiploids fell off drastically if the embryo resistance was more than 0.6 (Figure 2C) or if the relative fitness from somatic cleavage was less than 0.7 (Figure 2D). This was not the case for diploid drives, though such drives would of course be slowed in these circumstances. This is because diploids support male drive, which would be unaffected by either of these parameters, allowing the drive to continue increasing in frequency. If the drive can only increase in frequency in females, as in haplodiploids, it would lose all driving capacity if all drive carrying offspring of females also have resistance alleles caused by high embryo resistance allele formation or are unable to reproduce due to fitness costs from high somatic cleavage rates. We also examined the germline resistance allele formation rate, but it has little effect on the genetic load (Figure S2C–D).

**Figure 2.**
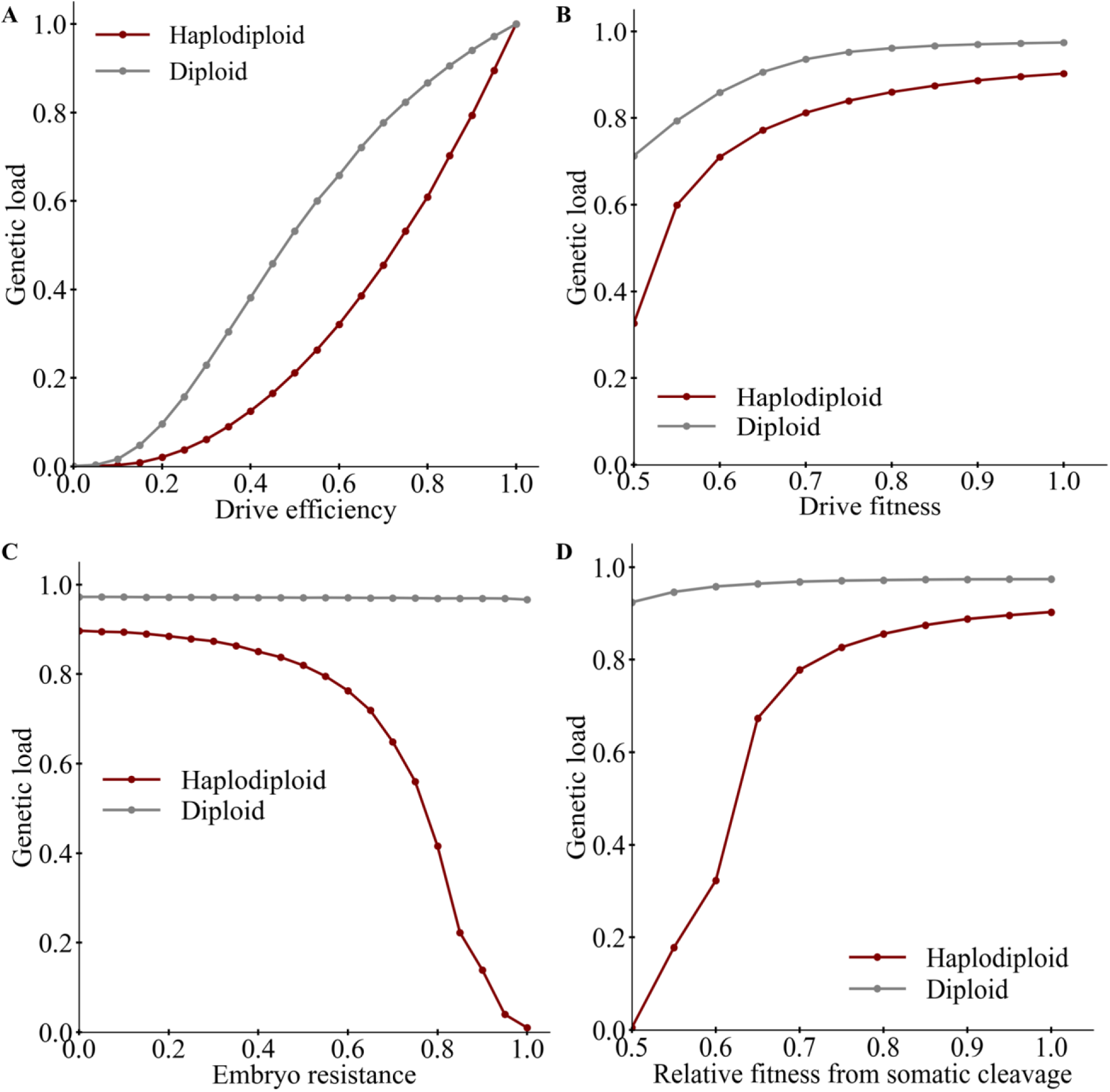
Effects of performance characteristics on the genetic load. The equilibrium genetic load is displayed for female fertility homing suppression drives with default performance characteristics and varying (**A**) drive conversion rate, (**B**) drive homozygote fitness, (**C**) embryo resistance allele formation rate, or (**D**) fitness cost in female heterozygotes due to somatic CRISPR cleavage activity. Displayed data is the average from 200 simulations.

### Drive performance in continuous space

We next examined drive performance in a model with dimensionless continuous space, where density dependence, mating, and offspring placement can now only occur locally. To focus on fundamental features of continuous space as related to drive, we again used a simple population genetic model in which local competition affects female fecundity. Previous studies indicated that suppression drives in continuous space could fail in several ways. In one, the genetic load would be insufficient to suppress the population, but because this also occurs in panmictic populations, we restricted our analysis to drives that would theoretically be able to suppress analogous panmictic populations. This included drives with at least 80% drive conversion rate and 80% fitness in homozygotes (with each drive allele contributing the square root of this fitness, and alleles having multiplicative fitness). A second mode of failure involves stochastic loss of the drive after the population size has been reduced due to the drive, making stochastic effects more dominant (early stochastic loss of the drive before it had a chance to spread never occurred in our model). This can also occur in panmictic populations, but it is far more common in models of suppression drives in continuous space^39,45^. This is because individuals carrying unique alleles at low population densities may struggle to find nearby mates, which doesn’t occur to nearly as great an extent in panmictic populations when all remaining individuals can potentially mate with each other. However, we found that this outcome was very rare for a female fertility homing suppression gene drive in haplodiploids, occurring 0.29% of simulations (Figure S3).

Another more common mode of drive failure was “chasing”. This refers to a phenomenon in which some wild-type individuals escape from the drive and reach empty regions where the drive had previously eliminated the population^39^. Here, they experience reduced competition and are able to have high numbers of offspring. The drive remains in contact with these wild-type groups, “chasing” them and continuously causing suppression, but in many cases, the drive fails to ultimately eliminate the population, even after long periods of time. Chasing was quite common in our simulations of haplodiploids. This was not unexpected, given similar results in other types of drives^39^ and given that our panmictic results that showed occasional rebound of the wild-type population (Figure 1B–C). Movies showing short and long chases involving the haplodiploid drive can be found on GitHub (https://github.com/jchamper/ChamperLab/tree/main/Haplodiploid-Suppression-Modeling).

Specifically, when drive fitness, and more importantly, drive efficiency (representing the drive conversion rate), were high, the population was often suppressed without chasing (Figure 3A, Figure S4A–B). However, it was more common for a period of chasing to occur prior to suppression (Figure 3B). When both efficiency and fitness were low, chasing could persist for extended periods of time, with the population still in a chasing state after 1000 generations (Figure 3C). With default parameters (Table S1), suppression could occur fairly quickly, though chasing could somewhat extend this (Figure S5). In simulations when chasing was avoided or minimized, suppression occurred quickly (Figure 3D). However, time to suppression (Figure 3D) was mostly controlled by the duration of chasing (Figure 3E) when chasing became significant at lower efficiency and fitness, sometimes lasting several hundred generations.

**Figure 3.**
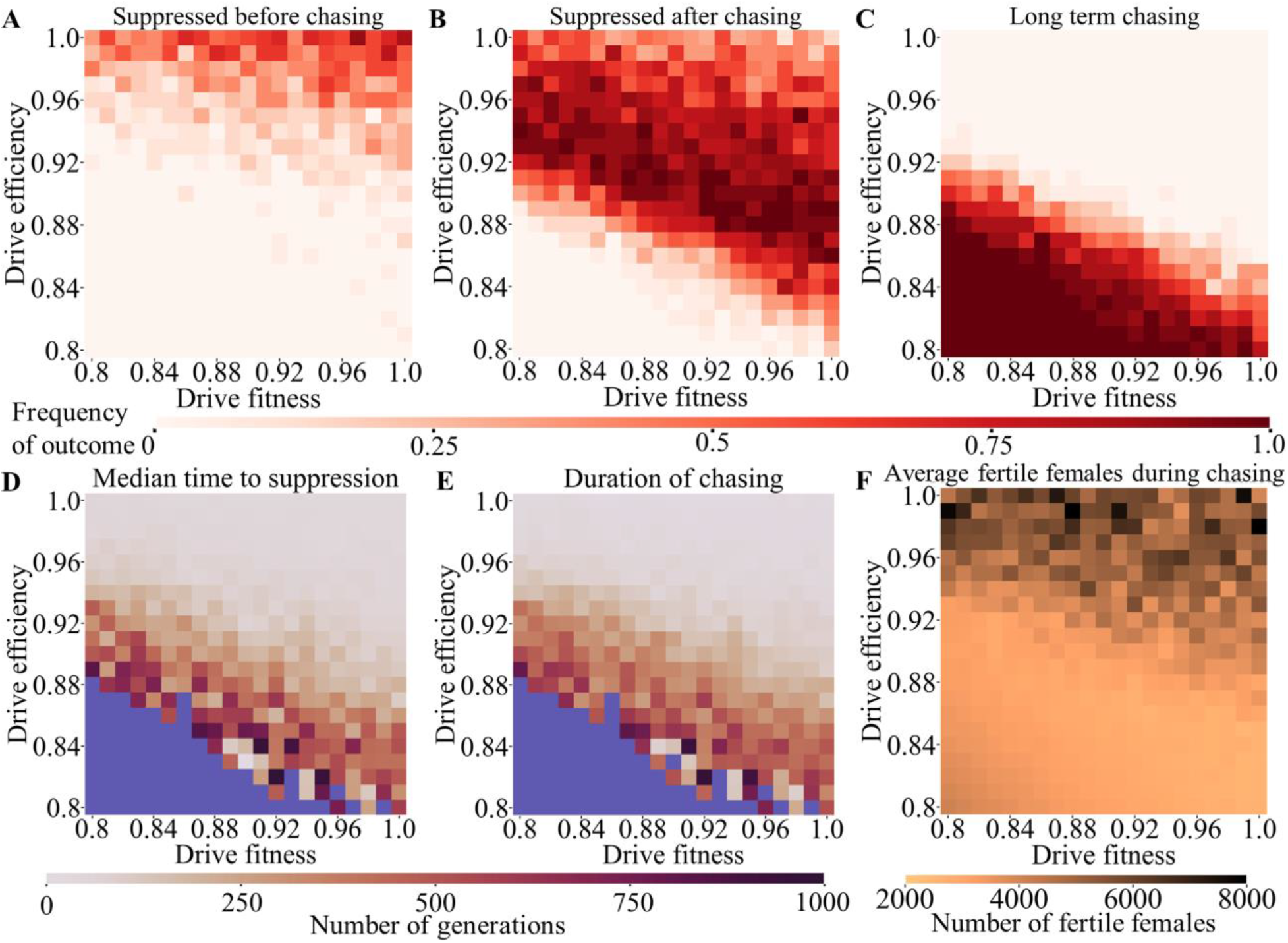
Outcomes of a drive release in continuous space with varying drive efficiency and fitness. Female drive heterozygotes and drive males with default parameters and varying efficiency and fitness were released in a into the middle of a spatial population of 50,000 individuals. Outcome rates are displayed for (**A**) suppression without chasing, (**B**) suppression after a period of chasing, and (**C**) simulations in which chasing was still occurring at the end of the simulation. Also displayed is (**D**) the median time to suppression, (**E**) the average duration of chasing that eventually ended in suppression (blue represents parameter space where chasing did not end for 1000 generations in any simulation), and (**F**) the average number of fertile females during any type of chasing. 20 simulations were assessed for each point in the parameter space.

In many cases, an outcome in which chasing persists could still provide benefits by reducing the population and its reproductive capacity. We measured this by tracking the average number of fertile females during a chase (Figure 3F). This indicated that even an inefficient drive could reduce the population several-fold compared to its initial state. This reduction was greater for more efficient drives. However, for drives with very high efficiency, the average number of fertile females tended to be higher with greater variance (Figure 3F). These were short-term situations that always led to complete suppression in short time scales (Figure 3B and 3D).

Ecological characteristics can also influence the outcome of a suppression drive release. We thus varied the migration rate and the population growth rate at low densities, similarly tracking drive outcomes for an efficient drive (Figure S4C–D). Unlike most previously investigated drives, we found that the low-density growth rate had little effect in the range tested (Figure 4, Figure S4D) (though eventually, high enough low-density growth rates would yield equilibrium outcomes in all situations, including panmictic populations). Migration rate had a similar effect as drive efficiency. When migration was high, suppression before chasing was common (Figure 4A), followed by suppression after chasing for most of the parameter range (Figure 4B). When very low, long-term chasing outcomes were common (Figure 4C). However, most chasing was short and quickly ended in suppression at middle and high migration rates (Figure 4D–E). When migration was low, the average population during a chase was higher (Figure 4F).

**Figure 4.**
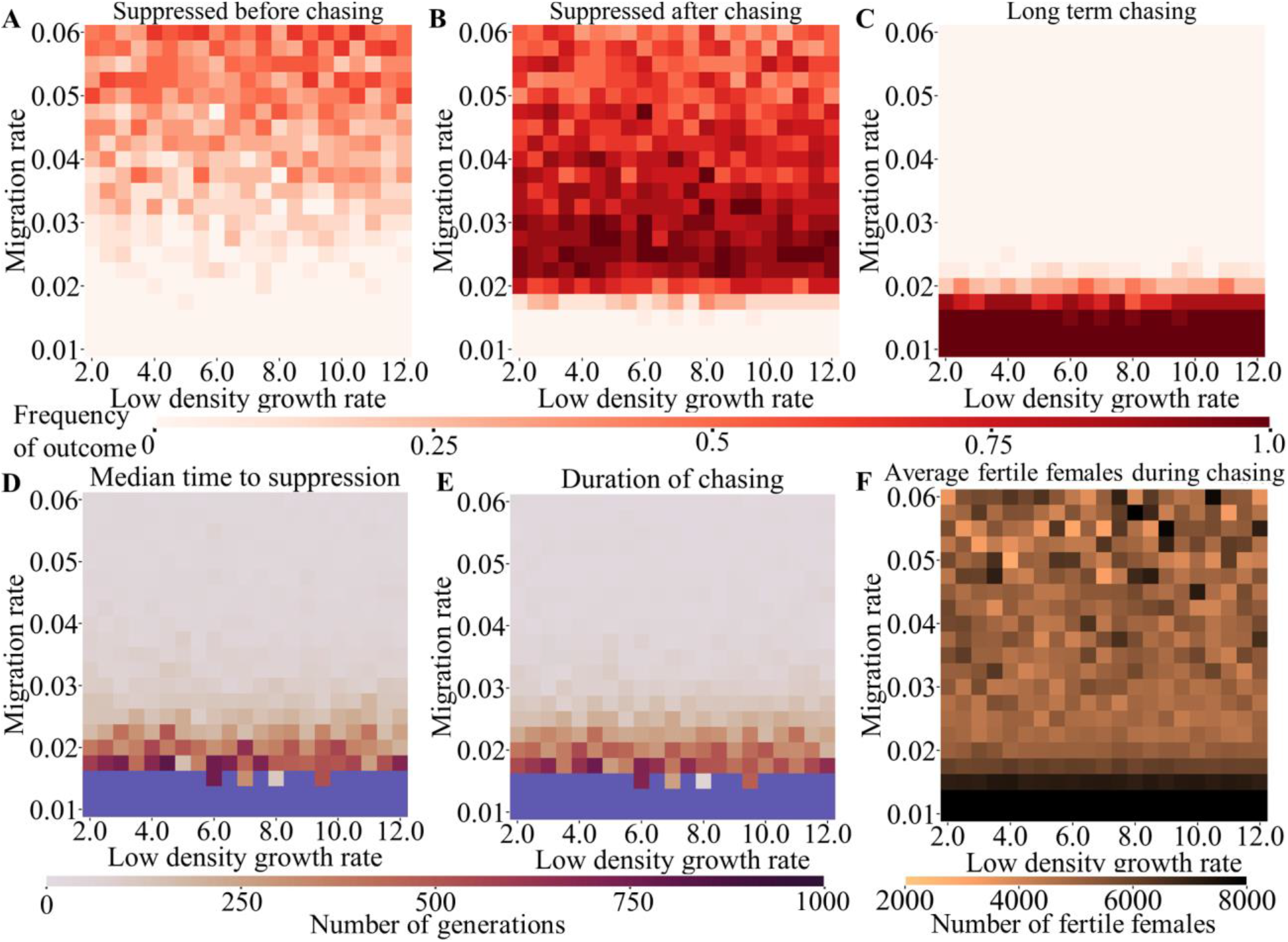
Outcomes of a drive release in continuous space with varying migration rate and low-density growth rate. Female drive heterozygotes and drive males with default performance parameters and varying migration and low-density growth rate were into the middle of a spatial population of 50,000 individuals. Outcome rates are displayed for (**A**) suppression without chasing, (**B**) suppression after a period of chasing, and (**C**) simulations in which chasing was still occurring at the end of the simulation. Also displayed is (**D**) the median time to suppression (**E**) the average duration of chasing that eventually ended in suppression (blue represents parameter space where chasing did not end for 1000 generations in any simulation), and (**G**) the average number of fertile females during any type of chasing. 20 simulations were assessed for each point in the parameter space.

We also examined the effect of population density on haplodiploid drives, which was previously shown to affect certain types of drive systems^39^. However, like homing suppression systems in diploids, we found that population density had little effect on outcomes, except for slightly increased frequency of chasing at very low population density (Figure S6).

Comparing haplodiploid and diploid drive performance in our spatial model, we surprisingly find that haplodiploid performance is only marginally worse (Figures S7–9, Table S2, and accompanying text). Haplodiploid drives suffer more from short-term chasing at high drive efficiency, but slightly less at lower efficiency and with lower rates of drive loss.

### Drive failure due to functional resistance allele formation

One critical issue that CRISPR homing suppression gene drives must overcome is that of functional resistance. If such alleles form, they are highly likely to quickly spread in the population and prevent successful suppression^16,23,25,26,28,39,41^. To assess functional resistance alleles in haplodiploids, we allowed each new resistance allele to have a small chance of being functional, which would then allow reproduction for any female with the functional resistance (r1) allele. The range of this chance spanned from 10^−6^, in which r1 alleles would almost never form in the simulations, up to 10^−3^, which could likely be achieved easily with even minimal effort toward finding conserved target sites or multiplexing with a single additional gRNA^16,43^. As with nonfunctional resistance alleles, they could not be converted to drive alleles. We expect haplodiploid drives to be more vulnerable to this, given the same rate of functional resistance allele formation. This is because haplodiploid drives have a 1:2 ratio for methods for drive increase (female germline) to methods of resistance allele formation (female germline and the embryos of drive females due to maternally deposited Cas9 and gRNA), while diploids have a higher 2:3 ratio (as above, but also drive increase and resistance allele formation in the male germline). This allows diploids to increase in frequency a greater amount for the same amount of resistance allele formation, thus allowing for more total resistance allele formation by the time the drive has reached high enough frequency to eliminate the wild-type population. Stochastic effects seen in Figure 1B–C can also increase the number of opportunities to form resistance alleles in haplodiploids compared to diploids. Yet, despite these factors, the rate of drive failure due to functional resistance allele formation was only modestly higher in haplodiploids in panmictic populations (Figure 5), consistent with previous studies of modification drives^30^. However, when drive conversion rate is reduced, haplodiploid drives become less able to suppress the population and thus more vulnerable to formation of functional resistance alleles (Figure S10 and accompanying text).

**Figure 5.**
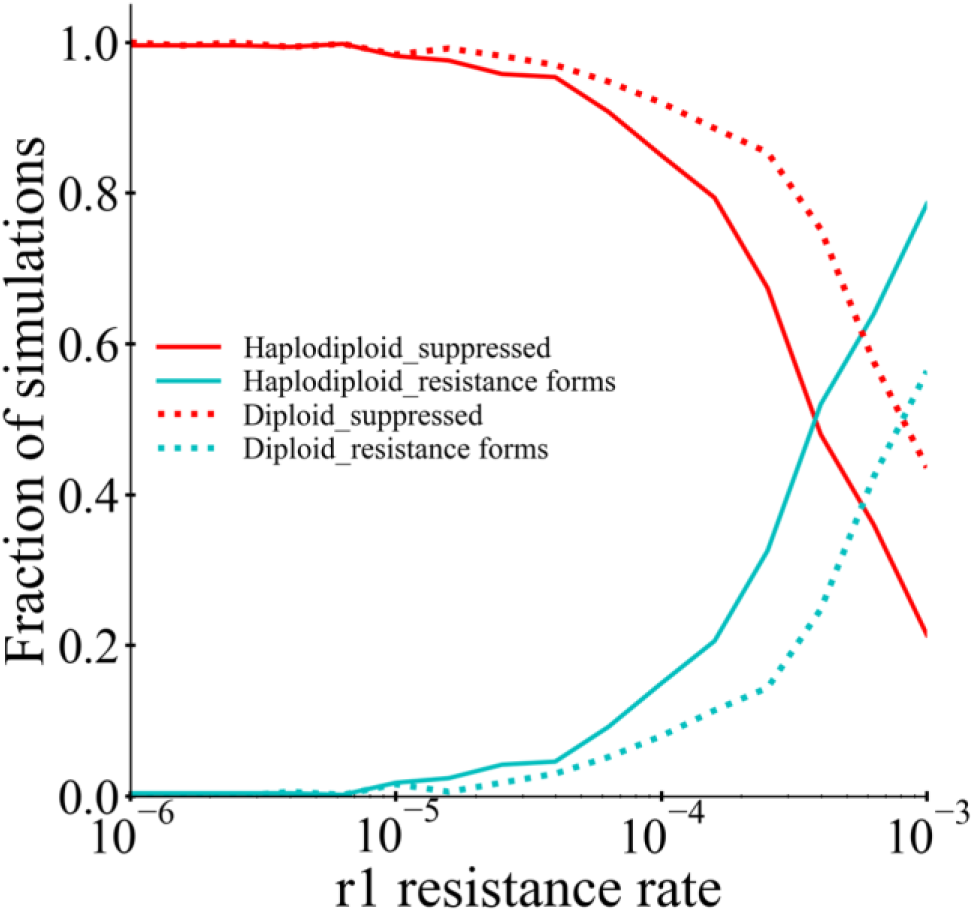
Functional resistance in panmictic populations. Drive heterozygotes (and drive males for the haplodiploid drive) with default performance characteristics and a variable relative rate of functional resistance allele (r1) formation were released at 1% frequency into a panmictic population of 50,000 wild-type individuals. The rate of successful suppression and drive failure due to resistance was tracked. 500 simulations were assessed for each point. Drive loss was not observed in any of these simulations.

In spatial populations, we expect that long periods of chasing would similarly increase the chance that functional resistance alleles could eventually develop, even if the rate of their formation would be at an acceptable level for panmictic populations in which suppression occurs quickly without chasing. This was previously demonstrated in diploid drives^39^. Examining our haplodiploid drives, we see this same phenomenon, with half of resistance outcomes taking place when resistance alleles formed after chasing with our default parameters (Figure S11). When we allow drive conversion rate and the relative rate of functional resistance allele formation to vary, a pattern emerges in which chasing substantially increases the parameter range over which functional resistance alleles are likely to form (Figure 6). Note that reductions in the drive conversion rate in these simulations do increase the absolute germline resistance allele formation rate (and therefore the functional resistance allele formation rate). This is generally consistent with the experimentally-derived notion that resistance allele formation represents a prevalent alternative to successful drive conversion^7,16,22,43^. Overall, we find that reducing the relative rate of functional resistance allele formation is important, but that it may be equally important to keep drive conversion high, both to reduce the absolute rate of resistance allele formation and to prevent opportunities to form resistance alleles during chasing.

**Figure 6.**
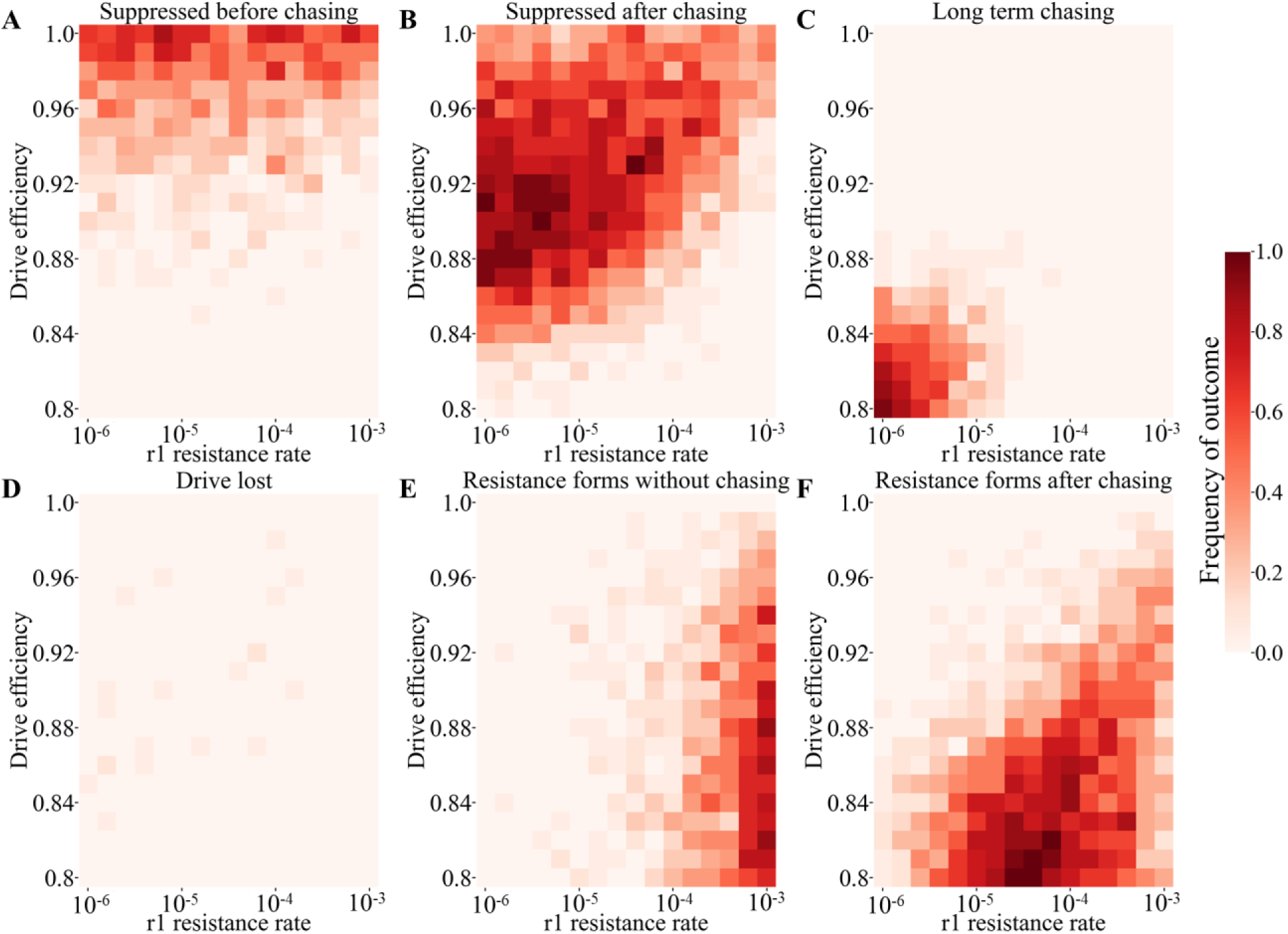
Functional resistance in spatial populations. Female drive heterozygotes and drive males with default performance parameters and varying drive efficiency and relative functional resistance allele formation rate were into the middle of a spatial population of 50,000 individuals. Outcome rates are displayed for (**A**) suppression without chasing, (**B**) suppression after a period of chasing, (**C**) simulations in which chasing was still occurring at the end of the simulation, (**D**) loss of the drive (this occurred before chasing in only one simulation), (**E**) drive failure due to formation of functional resistance alleles without chasing, and (**F**) drive failure due to formation of functional resistance alleles during chasing. 20 simulations were assessed for each point in the parameter space.

## Discussion

While modification type homing drives have previously been extensively analysed in haplodiploid organisms^30^, this study investigates the details of how such species could be suppressed by a gene drive. Our results show that this is possible in haplodiploid organisms, and that a highly efficient drive can actually provide for very strong and successful suppression. This remains true even in more challenging environments involving continuous space where the “chasing” phenomenon can slow or prevent suppression.

In haplodiploid organisms, the number of options for suppression gene drives is greatly reduced compared to diploids. Previously developed sex-biasing strategies based on sex chromosomes such as X-shredders are certainly not possible, and even drives that don’t rely on this will often lack the power needed to achieve high genetic loads. For example, homing drives targeting genes that are essential for both sexes or males would be eliminated quickly in males^34,35^, and even TADE^46,47^ drives would lack the power for suppression. Our modelling using the same format as in this study indicates that all of these configurations cannot increase in frequency on their own (TADE suppression slightly increases in frequency in the first generation only as the population equilibrates) and can only maintain a frequency in ideal form. Any imperfections in fitness or drive efficiency result in the drive declining in frequency regardless of release size. Modified versions of these homing drive strategies could still provide some suppression^34,35^. Fortunately, we found that the strategy of targeting a haplosufficient but essential female-specific gene with a homing drive was still able to achieve high genetic load (and though we modelled a female fertility gene, results would be broadly similar with a female viability gene). This type of gene drive has been well studied, with the latest drives even circumventing functional resistance alleles by careful target site selection^5^ or multiplexed gRNAs^42^. Because haplodiploid drives are somewhat more vulnerable to such functional resistance alleles, both methods would likely also be needed to avoid functional resistance in large, natural haplodiploid populations. Fortunately, though, such methods should function with similarly high efficiency to their demonstrations in flies and mosquitoes, meaning that sufficiently low resistance drives could likely be generated if baseline drive conversion efficiency of the target species was acceptably high. However, our study indicates that homing suppression drives in haplodiploids would need to keep somatic activity and embryo resistance allele formation even lower than in diploid drives to be effective. Though a rigorous requirement, there is still some flexibility in the specific values, and previously constructed drives have already achieved low enough somatic activity in flies^42^ and embryo resistance in mosquitoes^5^.

Surprisingly, we found that in continuous space models, female fertility homing suppression drives perform nearly as well in haplodiploid species as in diploids. Though more vulnerable to chasing over a fair amount of the parameter space we investigated, chases in most of this region were short, meaning that ultimate differences between the drives were minimal. Haplodiploid drives even had some minor advantages in some circumstances, such as reduced rates of drive loss and less loss of efficiency when the low-density growth rate was high. A previous study indicated that drives with less powerful mechanisms tended to be far more vulnerable to chasing^39^, but despite being unable to perform drive conversion in males to increase in frequency, this did not significantly apply to haplodiploid drives. This is perhaps because the drive can still diffuse quickly into wild-type populations but causes suppression more slowly than diploid drives, making it more difficult for pockets of wild-type individuals to stochastically avoid the drive in a chasing situation.

We note that our model is quite simplified compared to some realistic populations. We used discrete generations and simplified competition, while avoiding complex lifecycle characteristics that are typical of many species, especially many haplodiploid insects. In our spatial model, we only considered a bounded square and assumed spatial homogeneity. Our intention in this study was to present a basic overview of suppression drives in haplodiploids and how they differ from drives in diploids. However, this generality means that our conclusions are limited when considering specific species, particularly for predicting time intervals from a drive release to various outcomes. Lifecycles, diverse landscapes, connectivity between populations, types of intraspecific and interspecific competition (and other interactions), seasonality, and other factors can all influence how a drive may spread through and suppress a population.

Unlike TADE suppression drives, powerful homing drives are unconfined, even in haplodiploid organisms. This is because they can increase rapidly in frequency based on their drive allele conversion mechanism, even when initially present in a population at low frequency, allowing them to invade new populations with even small numbers of migrations. This may be undesirable in some situations such as targeted suppression of invasive populations. Tethered drive systems could provide a solution to this^48,49^ and allow for confined drive in haplodiploids. Indeed, modelling shows that a TARE drive, which could be used to confine a homing drive that lacks Cas9, would perform well at an X-linked locus^46^, and it would have identical performance in haplodiploid species in other configurations^47^ as well.

X-linked homing drives in diploids (which have been similarly neglected in previous modelling like haplodiploids) would also have similar performance to drives in haplodiploids^30^, and our modelling is thus equally applicable to both. Normally, we would expect autosomal diploid suppression drives to have equal or superior performance to X-linked drives. However, our modelling in continuous space indicates that there may be a narrow parameter space where an X-linked system would be preferred for avoiding long-term chasing outcomes. Of course, more realistic species-specific modelling and knowledge of model parameters would be necessary before development of an X-linked suppression drive could confidently be recommended over an autosomal version.

While development of gene drives in new species remains difficult, our results are nonetheless promising for future control of haplodiploid pests. Thus, studies developing CRISPR knock-in techniques in haplodiploid pest species, as well as characterization of germline promoter elements, female-specific target genes, and gRNA promoters can be considered high priority. As with diploids, ecological field investigations and more advanced modelling studies are also needed for accurate outcome predictions of gene drive deployment in specific species and regions of interest.

## Acknowledgements

Thanks to Samuel E. Champer and Isabel K. Kim for assistance with SLiM programming. This study was supported by laboratory startup funds from Peking University and the SLS-Qidong Innovation Fund.

## Supplemental Information

### Supplemental Methods

#### Modelling drive performance

For reproducing individuals in our model, the process of gene drive takes place independently in each gamete. The rate for wild-type alleles being converted into drive alleles in the germline of heterozygotes was determined by the “drive conversion rate”/“drive efficiency”. The probability of a wild-type allele instead being converted into a resistance allele in drive heterozygous females was equal to the “germline resistance allele formation rate”, which we set as 0.5*(1-drive conversion rate) in our model so that half of remaining wild-type alleles become resistance alleles after drive conversion takes place. If neither drive conversion nor resistance allele formation occurs, the wild-type allele is assumed to not be cut by the drive and will be transmitted to offspring normally. Embryo resistance was also taken into consideration, with a variable “embryo resistance allele formation rate” in which wild-type alleles of new offspring (which could come from either the mother or the father) are converted to resistance alleles if their mother has at least one drive allele (this rate is specified for drive heterozygotes and is somewhat higher and lower for drive homozygotes and drive/resistance allele heterozygotes, respective, according to our previous model^16^). If the offspring’s mother possesses a gene drive allele while the offspring has any remaining wild-type target sites, they might be converted into resistance allele due to persistence of maternally deposited Cas9 and gRNA in the embryo. The “relative r1 formation rate” determines the chance that any new resistance allele will be a functional/r1 allele, preventing sterility in any female that is a carrier of such an allele. Our default parameters (Table S1) represented an efficient drive^5,8,9^, with 95% drive conversion rate and a 5% embryo resistance allele formation rate. These represent what has been achieved in Anopheles mosquitoes, and the relative ly efficient default parameter also allows us to investigate other variables, showing which could potentially cause a drive to fail and in general allowing our simulations to have dynamic outcome ranges.

Drives could also have reduced fitness from two sources. Direct fitness effects were specified by the main fitness parameter, which previously referred to the fitness relative to wild-type in drive homozygotes, with multiplicative fitness per drive allele. For haplodiploids, this parameter is directly used for males with one drive allele. It affects female fecundity directly, and fitness also proportionally reduces the chance that a male will be selected as a mate. Our default fitness value was 0.95, indicating a high quality but imperfect drive^50^. Negative fitness effects can also stem from somatic expression in female drive/wild-type heterozygotes, which can disrupt wild-type alleles in somatic cells that are necessary for female reproduction^5,27^. Our default for this parameter is 1, indicating no effects on female fitness, but we also investigated effects of varying this parameter from 0.5 to 1, where it has the effect of multiplying the fecundity of drive/wild-type heterozygous females. Aside from these considerations, note that females with only drive and/or nonfunctional resistance alleles will always be completely sterile due to the target gene of the drive, having an effective fitness of zero.

#### Panmictic population model

To determine if gene drive is feasible for haplodiploid suppression, we first implemented the drive in a panmictic population. The simulation is initialized with a single release of gene drive heterozygous females and drive carrying males at a frequency of 1% of the total population after allowing the population to equilibrate for 10 generations. Fertile females could pick a male to mate with at random, and then the equation *ω*’_*i*_ = *ω*_*i*_\**β* / [(*β*−1) N/K+1] is used to calculate the female fecundity, where *ω*_*i*_ is the baseline fecundity due to genotype, N is total population size, K is the population carrying capacity, and *β* is the low-density growth rate (the fitness advantage individuals experience when unaffected by competition due to lack of close neighbours). Changing this parameter would not substantially affect drive frequencies trajectories in panmictic populations, except in the final stages where it could eventually prevent suppression if very high. However, it has previously been shown to more strongly affect outcomes in spatial scenarios^16^. We use a default value of 6 for the low-density growth rate, in keeping with our previous study^16^ and allowing rapid growth at low densities, but also allowing the efficient drives we consider here to suppress the population in favourable conditions. We assumed the number of offspring conform to a binomial distribution and then draw 50 independent offspring with survival *ρ* = *ω*_*i*_’/25. This means that a wild-type female will produce two offspring on average if the population is near its capacity. We note that in this discrete-generation model, competition affects female fecundity. In some cases, this could be an accurate representation of a particular species. In in others, it represents an approximation of viability-based larval competition, where multiple adults depositing eggs in the same area would have fewer surviving offspring due to intense competition at the larval or juvenile level.

#### Determination of the genetic load

The genetic load is one way to assess the reproductive burden the drive places on the population. It refers to the fractional reduction in population reproductive capacity compared to a similar wild-type population, thus measuring the strength with which a gene drive can suppress the population. It is equal to 1 - (actual population during the next time step) / (expected next generation population if all individuals were wild-type). To allow for accurate measurements of the genetic load when it is high, population suppression was artificially prevented by allowing fertile females to have more offspring than normal during simulations used to measure genetic load^47^. Specifically, when enabled, additional offspring are generated for all fertile females increasing their number of offspring by a factor of 1/(fraction of females that are fertile). This allows the population to persist even if it would otherwise be reduced and eliminated. This was necessary when the genetic load of the drive was high enough to eliminate the population, preventing extended measurements when the drive was at equilibrium. In all genetic load simulations, the release size of the gene drive heterozygotes was 20% of the total population, referring to the number of drive-carrying individuals introduced to the wild-type population at the beginning of the simulation. The simulation was allowed to run for 50 generations for the drive to reach its equilibrium frequency, and then the last 100 generations of the simulation were considered for calculation of the average genetic load. The total population size was also adjusted to 100,000 to minimize stochastic effects and increase the accuracy of the genetic load determination.

#### Spatial model

To conduct a more realistic assessment of drive performance, the panmictic model was expanded to a model involving continuous space. The 50,000 initial wild-type individuals were confined into a 1 x 1 arena, and the released drive-carrying individuals are all placed in a circle of radius 0.01 at the centre of the arena. Because our model is generic, these distances are necessarily unitless, so it would vary between species and perhaps locations in any specific scenario. Depending on the species and local ecology, it could potentially represent a wide area many kilometres long with many large ant colonies, or a small area, with slow-moving reproducing individuals dispersing in a small town of just several hundred meters. We initialized the simulation by releasing 500 female drive heterozygotes and drive carrying males (250 each, representing 1% of the total population) into the centre after allowing the simulation to equilibrate for 10 generations.

In the spatial model, the females were limited to choosing mates within a radius equal to the *migration rate* parameter (with a default of 0.04 to show a dynamic range of outcomes when drive efficiency parameters vary), and the females would not be able to reproduce if there wasn’t a male to mate with around that circle. To model local competition, we scaled the fecundity of females to *ω*’_*i*_ = *ω*_*i*_\**β*/[(*β*−1) *ρ*_*i*_/*ρ*+1], where *ρ*_*i*_ represented the number of other individuals around a female within a circle of radius 0.01, and *ρ* was the carrying density (equal to K/total area). Due to an abundance of resources and reduced competition, this allows a female to have more reproduction if she was in a low-density area. Offspring were distributed randomly around the mother at a distance drawn from a normal distribution with zero mean and standard deviation equal to the migration rate. The result of this process will produce an average displacement equal to *migration rate* 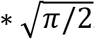. Because most species compete at smaller distances (often at larval or colony stages with low mobility compared to flying adults) than they can disperse, we fixed the migration distance to be equal or greater to the competition distance. The maximum migration value was set to show a range of outcomes and to allow drive “waves of advance” to form in the centre before any drive individuals reached the edge of the simulation arena. If the offspring was placed outside of the arena, their position is redrawn until they fall within the boundaries. This results in a slightly increased density near the boundaries of our simulation arena. Simulations are stopped 1,000 generations after the drive release if the population or the drive allele was not eliminated earlier.

#### Assessment of the “chasing” phenomenon

To quantify the degree of clustering of the population in the spatial model, we calculated Green’s coefficient (*G*) of each generation through the equation 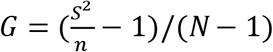, where *N* was the total population, *n* represented the mean number of the individuals and *s*^*2*^ was the variance of the counts. For this purpose, we divided the arena into an 8 x 8 grid and counted the wild-type homozygotes of each square.

In suppression drive, we often see the “chasing” phenomenon between wild-type and drive individuals after releasing drives into the population^16^. The population around the centre where the suppression drive is released could be eliminated rapidly, but then recolonized by escaping wild-type individuals. At the peak of this process, population is low, but Green’s coefficient is increased as well^16^. The escaped wild-type individuals produce more offspring because of the lack of competition, and soon they occupy more space, decreasing Green’s coefficient in response while increasing total population. The drive is still present, however, resulting in cycles of suppression and recolonization that have the appearance of the drive “chasing” wild-type across the landscape. Based on previous assessment^16^, we consider a chase to have begun in the lower generation of the first local maximum in Green’s coefficient and the first local minimum in the number of wild-type individuals. It is noted that we only count the wild-type homozygote individuals in the estimation since this will more closely correlate with chasing^16^.

#### Data generation

Simulations were performed on the High-Performance Computing Platform of the Center for Life Science at Peking University. Python was used to analyse the data and prepare the figures. The SLiM script, Python files, raw data, and chasing movies are availab le on GitHub (https://github.com/jchamper/ChamperLab/tree/main/Haplodiploid-Suppression-Modeling).

**Table S1.** Default Parameters.

| Parameter | Value |
| --- | --- |
| Drive conversation rate | 0.95 |
| Germline resistance formation rate | $0.5 \times (1 - \text{Drive conversion}) = 0.025$ |
| Embryo resistance formation rate | 0.05 |
| Homozygote fitness value | 0.95 |
| Female fitness from somatic activity | 1 |
| Low-density growth rate | 6 |
| Population carrying capacity | 50,000 |
| Drive release fraction | 0.01 |
| Drive release radius | 0.01 |
| Density interaction radius | 0.01 |
| Migration rate | 0.04 |
| Spatial arena size | 1 x 1 |

All models use the listed default parameters unless otherwise specified.

### Supplemental Results

#### Effect of performance parameters on the rate of drive spread

To examine how drive performance parameters affect the rate at which the drives spread, we measured the haplodiploid drive allele frequency 16 generations after the drive individuals were released, which we found to give a wide spread of values, making it suitable for comparing drives with different performance (at lower generations, drive frequencies would often not have time to increase much, while at higher generations, all drives would have reached high frequency, making comparisons more difficult). As expected, the drive conversion rate (the drive efficiency) and fitness were critical for rapid spread of the drive (Figure S1A). The germline resistance scarcely affected the suppression drive (Figure S1B), though it could only vary it from 0 to 0.05 due to high default drive efficiency. The drive allele frequency had a similar downward trend from increasing the embryo resistance allele formation rate (such alleles form due to maternally deposited Cas9 and gRNA in the embryo) and the relative fitness from reduced female fertility due to somatic cleavage (Figure S1). This is because large reductions in the reproductive capacity of females (due to somatic cleavage eliminating wild-type alleles in some cells, which are required for fertility) or their offspring (due to embryo resistance alleles resulting in sterile daughters) can directly counterbalance drive conversion in females, preventing the drive from being able to increase in frequency.

**Figure S1.**
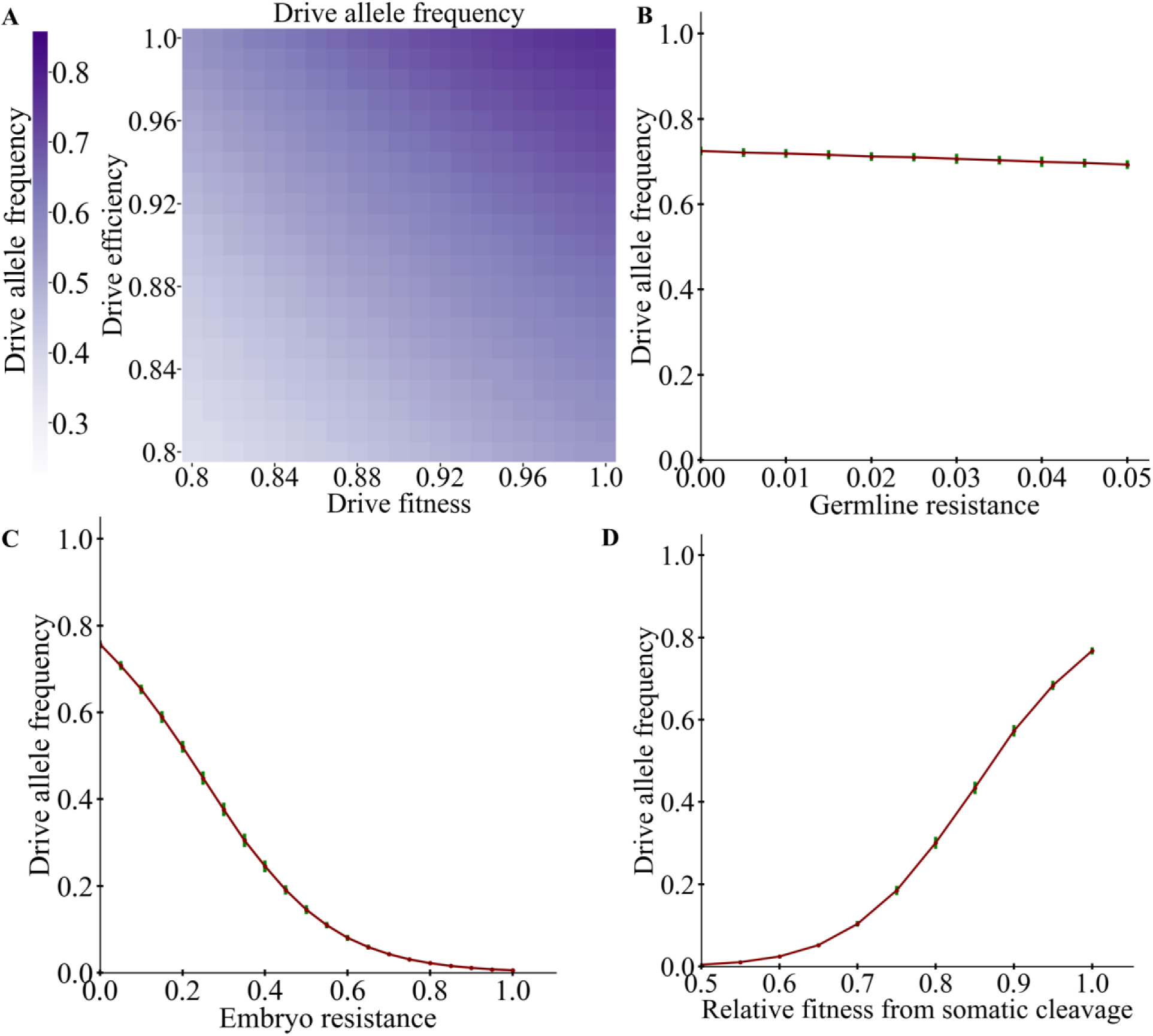
Effects of performance characteristics on haplodiploid drive frequency at generation 16. Drive heterozygotes (and drive males for the haplodiploid drive) with default parameters were released at 1% frequency into a panmictic population of wild-type individuals. The drive allele frequency is displayed 16 generations later for varying (**A**) efficiency and homozygote fitness, (**B**) germline resistance allele formation rate, (**C**) embryo resistance allele formation rate, or (**D**) fitness cost in female heterozygotes due to somatic CRISPR cleavage activity. Displayed data is the average from 20 (**A**) or 200 (**B-D**) simulations. Error bars represent the standard deviation.

**Figure S2.**
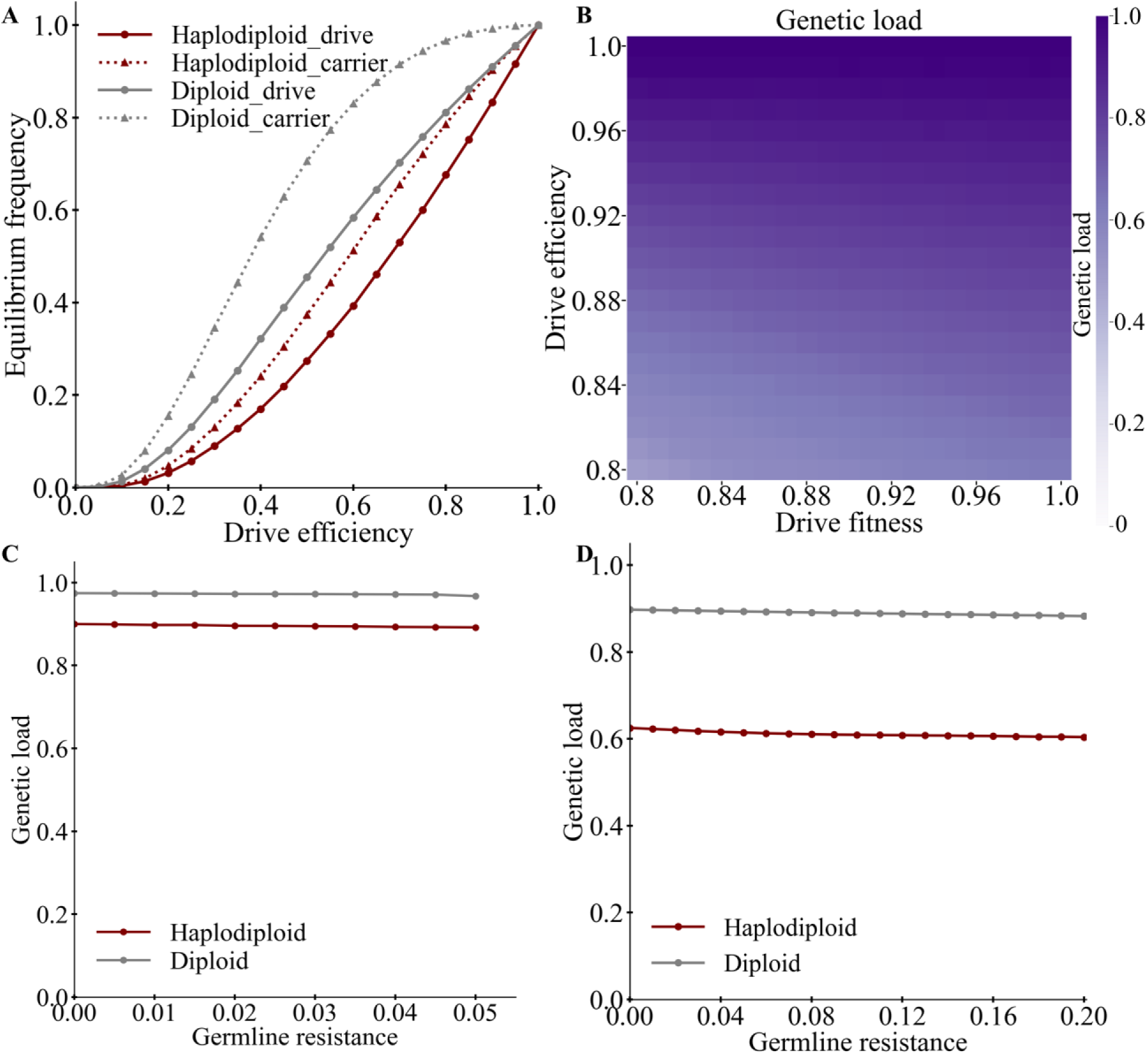
Effects of additional performance characteristics on the genetic load and drive equilibrium frequencies. (**A**) The drive allele and carrier frequencies are displayed for both haplodiploid and diploid drives at equilibrium when maximum genetic load has been reached. The genetic load for female fertility homing suppression drives at equilibrium is displayed for drives with default performance characteristics except for (**B**) varying drive efficiency and homozygote fitness or (**C**) varying germline resistance allele formation rate. We additionally (**D**) varied germline resistance rate while setting the drive conversion rate to 80%. Displayed data is the average from (**A-B**) 20 or (**C-D**) 200 simulations per data point.

**Figure S3.**
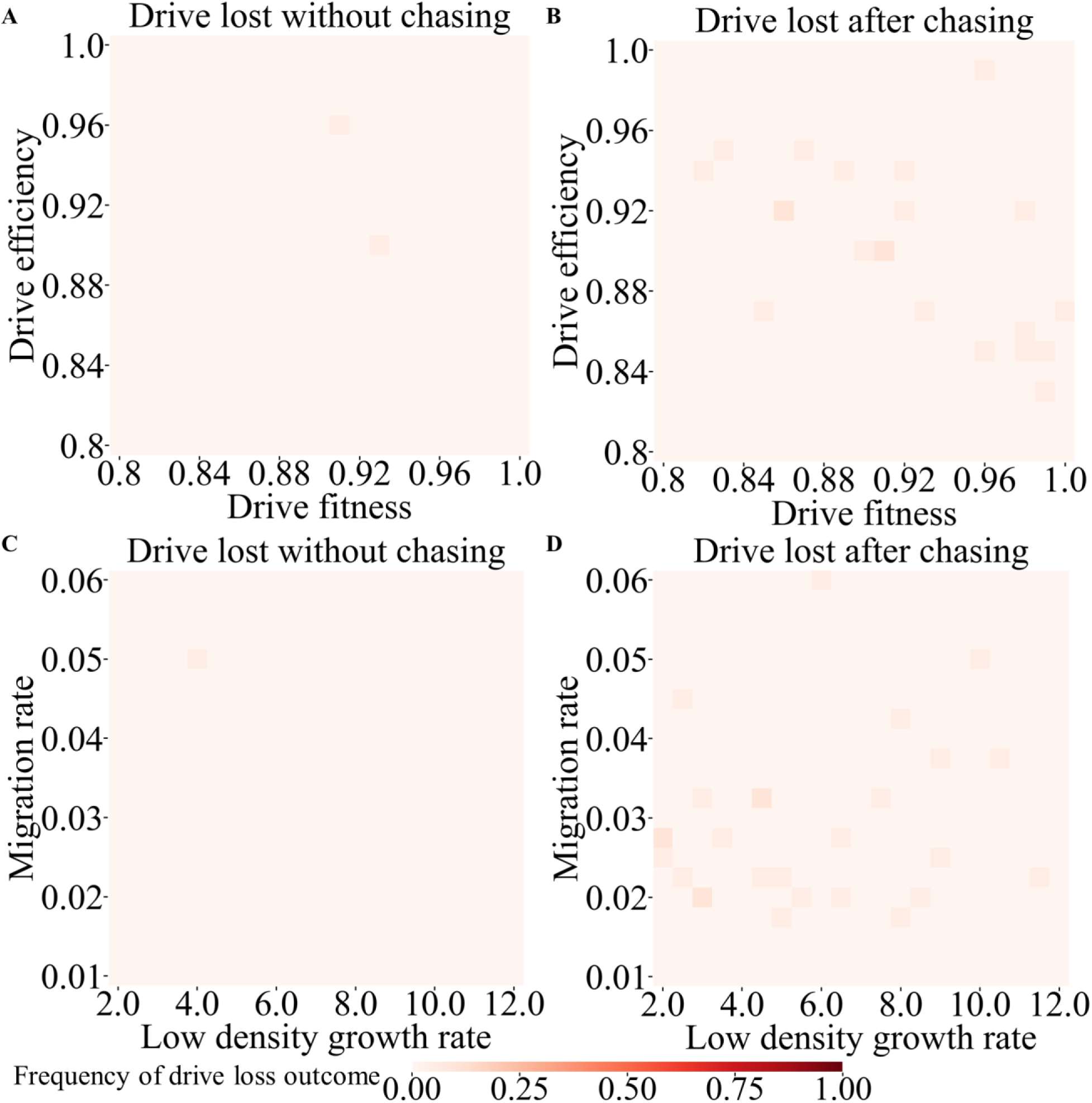
Drive loss outcomes in continuous space. Female drive heterozygotes and drive males were released into the middle of a spatial population. Default parameters were used with (**A,B**) varying drive efficiency and fitness and (**C,D**) varying migration rate and low-density growth rate. Rates of drive loss (**A,C**) without chasing or (**B,D**) after chasing are displayed. 20 simulations were assessed for each point in the parameter space.

**Figure S4.**
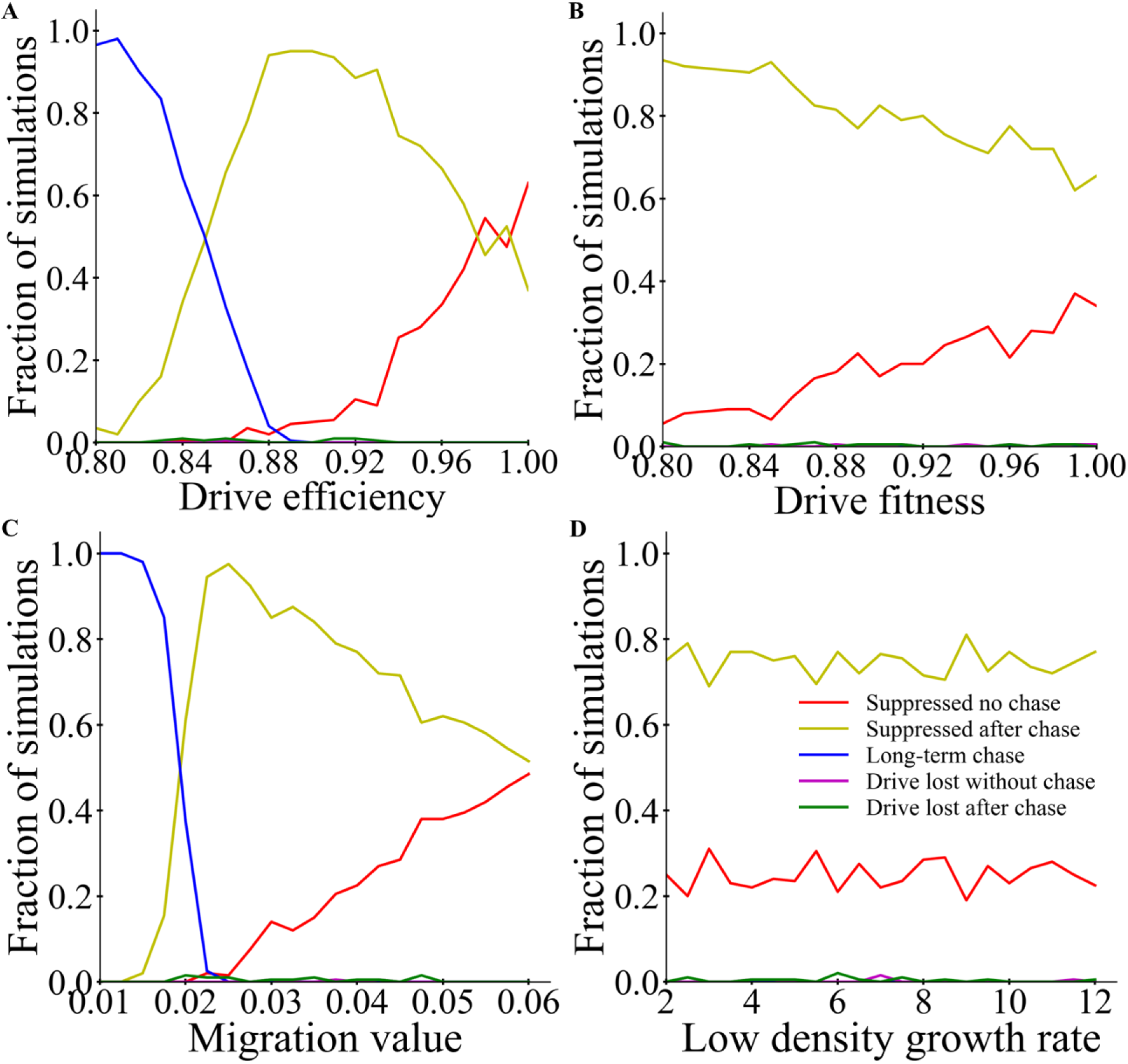
Outcomes of a drive release in continuous space. Female drive heterozygotes and drive males with default parameters and varying (**A**) efficiency, (**B**) fitness, (**C**) migration rate, and (**D**) low-density growth rate were released into the middle of a spatial population. The rate of each outcome type is displayed. 200 simulations were assessed for each point.

**Figure S5.**
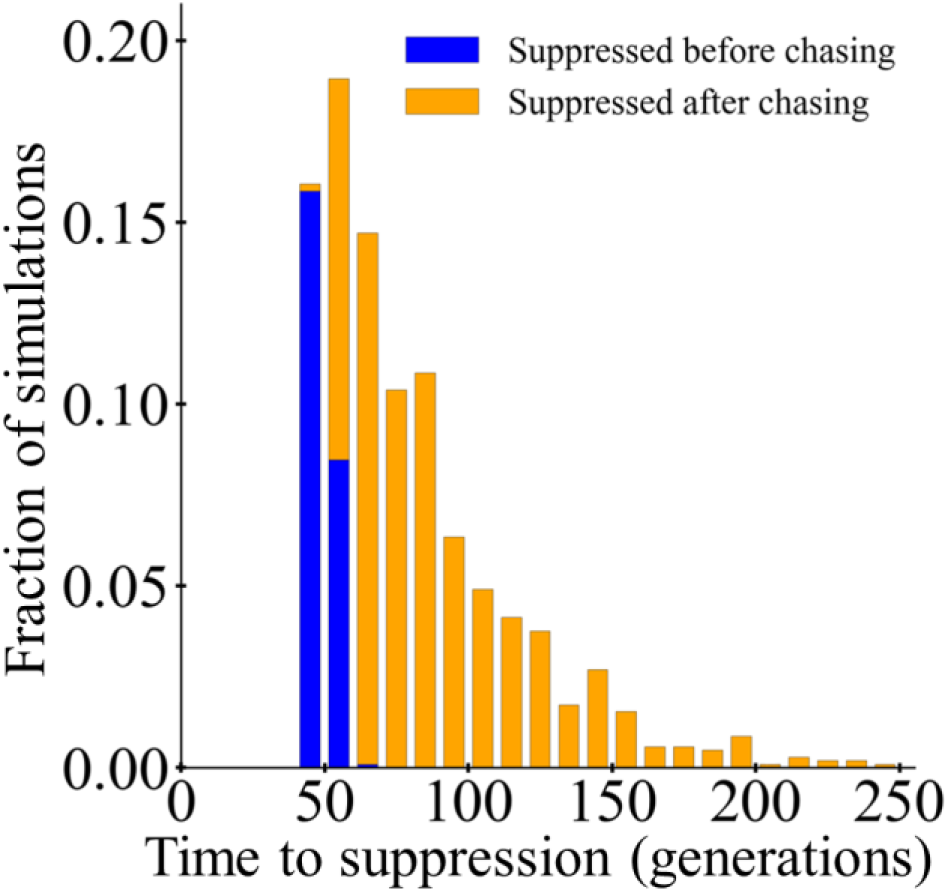
Outcomes of a drive release in continuous space. Female drive heterozygotes and drive males with default parameters were released into the middle of a spatial population. The time to population elimination is shown for each simulation, as well as whether chasing occurred. 1040 simulations were assessed, of which 1034 resulted in suppression and are displayed. In one simulation, suppression did not occur due to long-term chasing, and in five simulations, the drive was lost after a period of chasing.

**Figure S6.**
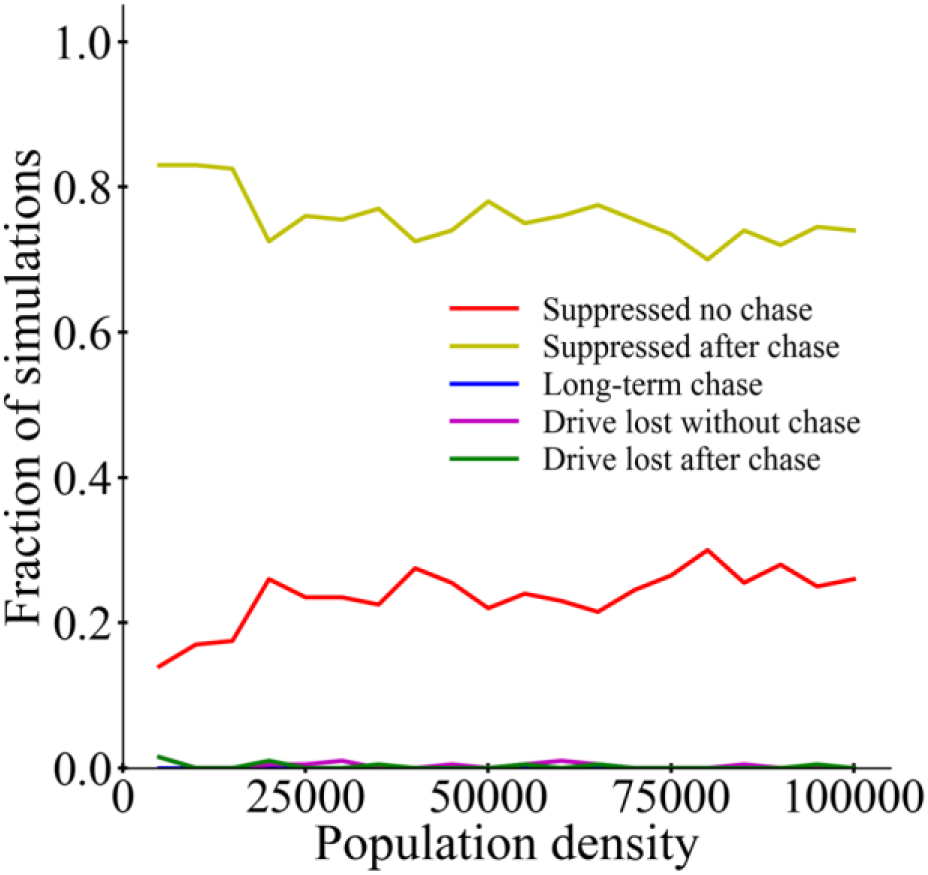
Effect of population density on drive outcomes in continuous space. Female drive heterozygotes and drive males with default parameters were released into the middle of a spatial population. The rate of each outcome type is displayed for varying population density. 200 simulations were assessed for each point.

#### Comparison between haplodiploid and diploid drives in the spatial model

Our previous study showed that in diploids, a female fertility homing suppression drive was also vulnerable to the chasing phenomenon in continuous space. However, it was still considered to be a good quality drive and performed better that homing suppression drives targeting genes that would cause sterility or nonviability in both sexes, as well as driving Y chromosomes (based on X-shredders)^39^. To measure how such a diploid drive compared to our haplodiploid drive, we compared their performance in identical conditions (Figure S7–8) with special attention to how performance varies with drive conversion efficiency (Figure S9). Despite the substantially higher genetic load and rate of increase of the diploid drive, both drives had similar performance in terms of final outcomes. Over a wide range of parameter space, the haplodiploid drive tended to have outcomes where chasing was followed by suppression (Figure S7B, S8B) when suppression would occur without chasing in the diploid drive (Figure S7A, S8A). However, in most of this range, the chases were very short (Figure 3D, 4D, S9), thus causing only small delays before complete suppression. In other areas of parameter space, the haplodiploid drive actually had slightly lower rates of chasing, especially when the drive was less efficient (Figure S7C, S9) or when the low-density growth rate was high (Figure S8C). Indeed, the haplodiploid drive in general was not substantially negatively affected by higher low-density growth rates, in contrast to other drives^39^ (though if this parameter becomes high enough, than any drive with a genetic load of less than 1 would eventually lose its ability to suppress the population and instead reach an equilibrium state). Also of note, the haplodiploid drive experienced drive loss much less than the diploid drive (Figure S7D–E, S8D–E, S9). This is possibly because the diploid drive, which is faster, can cause local elimination before the drive spreads to adjacent areas in some cases. The haplodiploid drive, while also capable of causing local elimination, does this more slowly, allowing the drive to disperse into adjacent wild-type areas before it is lost.

To assess the reasons for the female fertility homing suppression drive’s similar performance in haplodiploids and diploids despite differences in genetic load and rate of increase, we assessed panmictic population properties when the drive had reached 90% frequency with otherwise default parameters^39^ (see Table S2 and accompanying text). This is analogous to the situation on one side of the drive’s “wave of advance” where the population is nearly fully suppressed^39^. We found that the haplodiploid drive has a higher population of fertile females at 90% frequency compared to the diploid drive, thus reducing the chance that chasing can be initiated by stochastic escape of wild-type individuals from this region. This advantage can explain why the same drive in haplodiploids and diploids generally had similar performance, despite the greater power of the drive in diploids.

**Figure S7.**
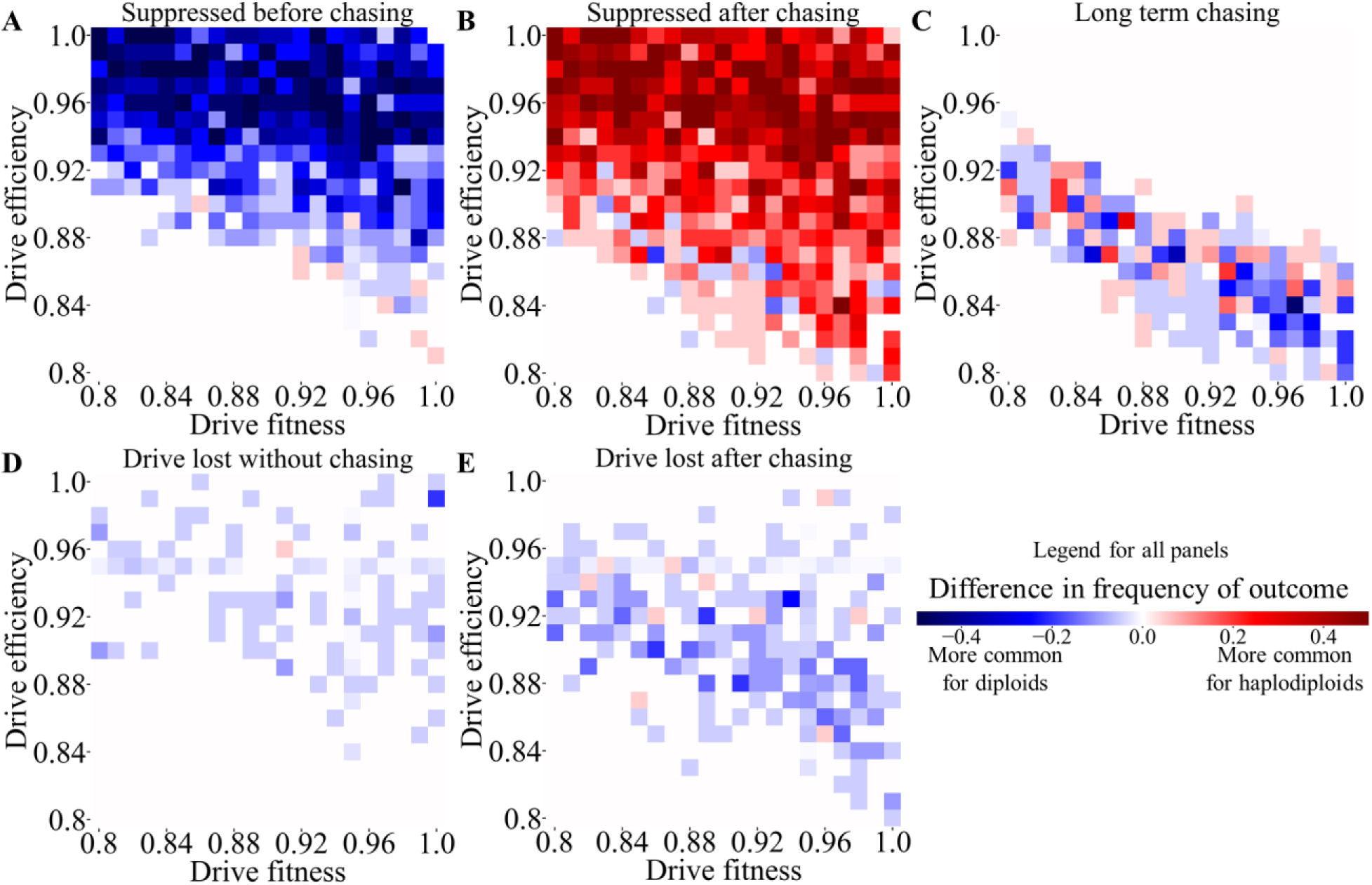
Comparison of continuous space outcomes between haplodiploids and diploids with varying drive efficiency and fitness. Drive heterozygotes (and drive males for the haplodiploid drive) with default parameters and varying drive efficiency and fitness were released into the middle of a spatial population. Differences in outcome rates between haplodiploids and diploids are displayed for (**A**) rapid suppression without chasing, (**B**) suppression after a period of chasing, (**C**) simulations in which chasing was still occurring at the end of the simulation, (**D**) drive loss without chasing, and (**E**) drive loss after a period of chasing. Red means that the outcome occurs more often in haplodiploids, and blue for diploids. The outcome itself could have occurred at high or low absolute rates in either. Only the difference is displayed. 20 simulations were assessed for each point in the parameter space. Data for diploid individuals is from a previous study^39^.

**Figure S8.**
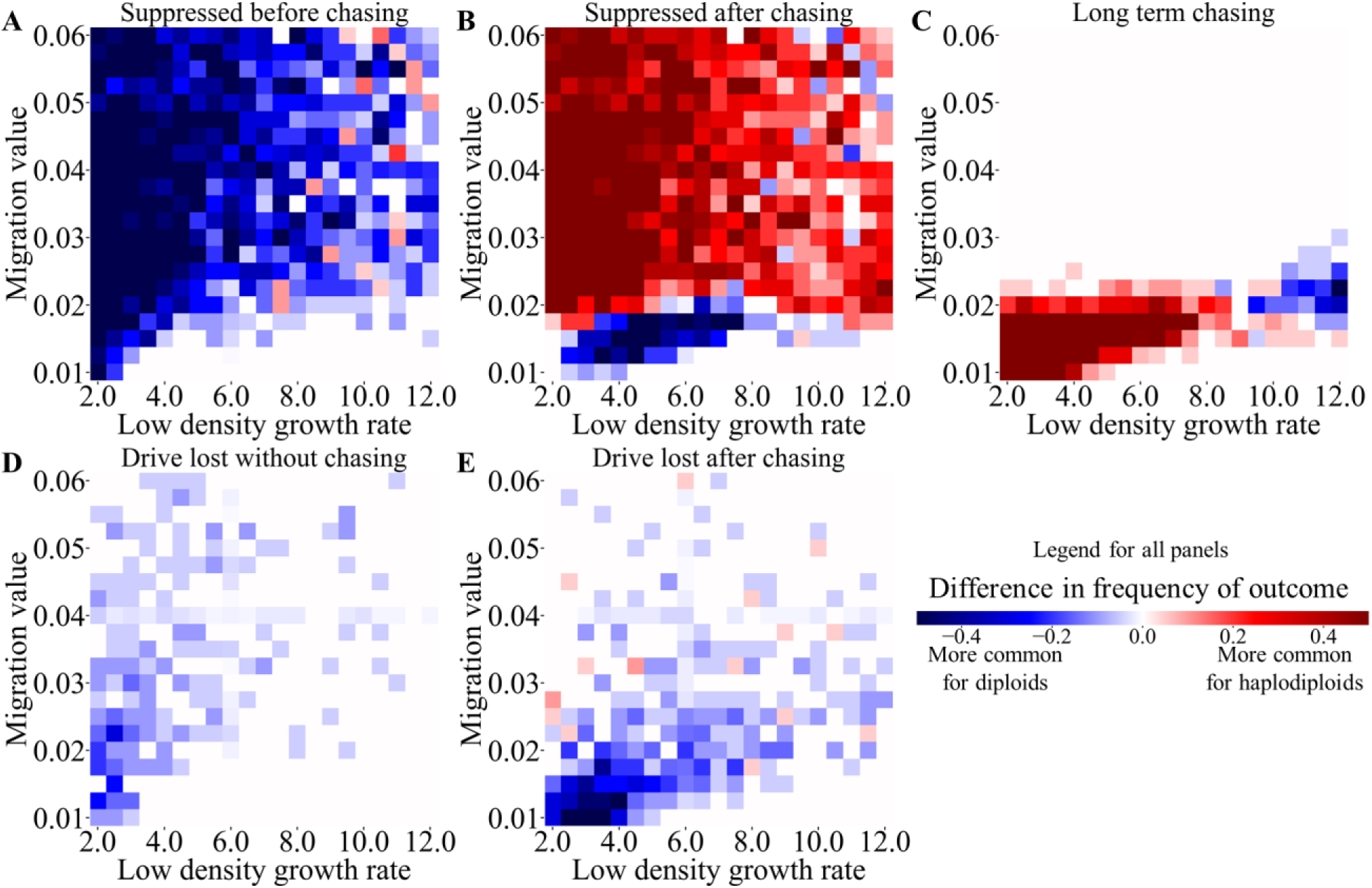
Comparison of continuous space outcomes between haplodiploids and diploids with varying migration and low-density growth rate. Drive heterozygotes (and drive males for the haplodiploid drive) with default drive performance parameters and varying migration and low-density growth rate were released into the middle of a spatial population. Differences in outcome rates between haplodiploids and diploids are displayed for (**A**) rapid suppression without chasing, (**B**) suppression after a period of chasing, (**C**) simulations in which chasing was still occurring at the end of the simulation, (**D**) drive loss without chasing, and (**E**) drive loss after a period of chasing. Red means that the outcome occurs more often in haplodiploids, and blue for diploids. The outcome itself could have occurred at high or low absolute rates in either. Only the difference is displayed. 20 simulations were assessed for each point in the parameter space. Data for diploid individuals is from a previous study^39^.

**Figure S9.**
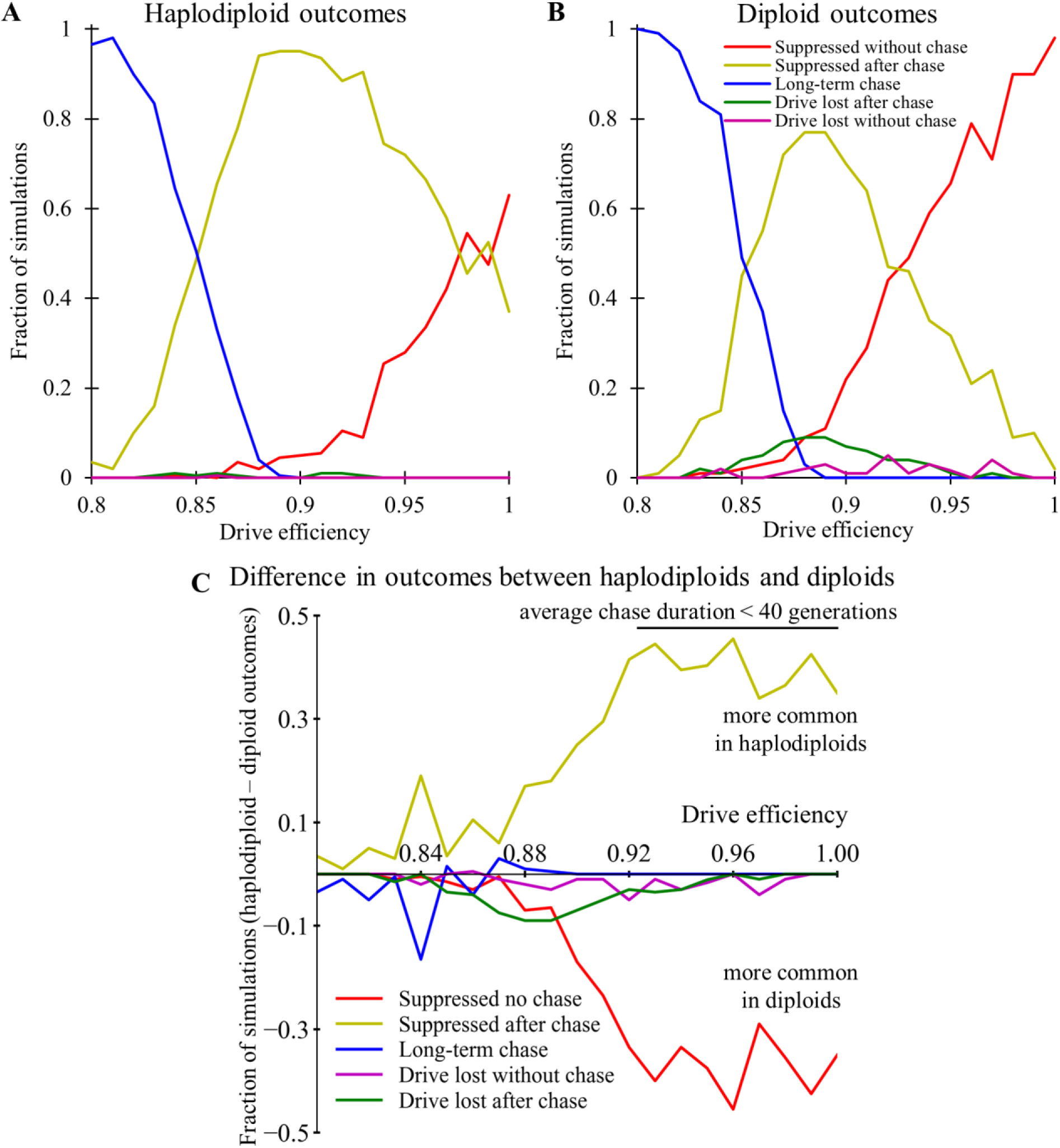
Comparison of outcomes in continuous space between haplodiploid and diploid suppression drives. Drive heterozygotes and drive males for the (**A**) haplodiploid drive and drive heterozygotes for the (**B**) diploid drive with default parameters and varying drive efficiency were released into the middle of a spatial population, and outcomes were tracks. (**C**) The differences in the outcome rates between haplodiploids and diploids is displayed (higher values represent outcomes that are more common in haplodiploids, while lower values correspond to outcomes that are more common in diploids). 200 simulations were assessed for each point. Data for the drive in diploids is from a previous study^39^.

#### Functional resistance in panmictic populations with varying drive conversion

When drive conversion is low, haplodiploid drives are considerably more sensitive to functional resistance alleles, with suppression failing over a wider range of parameters (Figure S10A), usually due to formation of resistance alleles (Figure S10B). This is because low drive conversion can greatly delay suppression due to low genetic load being unable to overcome the increased growth rate at low density. In this case, suppression can still occur due to stochastic fluctuation, but this extends the time during which a functional resistance allele can form. If drive conversion is sufficiently low, then suppression can also fail due to the drive and wild-type alleles remaining in equilibrium, with even normal levels of stochastic fluctuation unable to induce suppression for long periods of time (Figure S10C). Note that we did not see this “equilibrium” outcome in any of our spatial populations in this study due to lower local population sizes and thus greater stochasticity (such equilibrium outcomes are possible in spatial populations for other types of drives with sufficiently low genetic load^39^).

**Figure S10.**
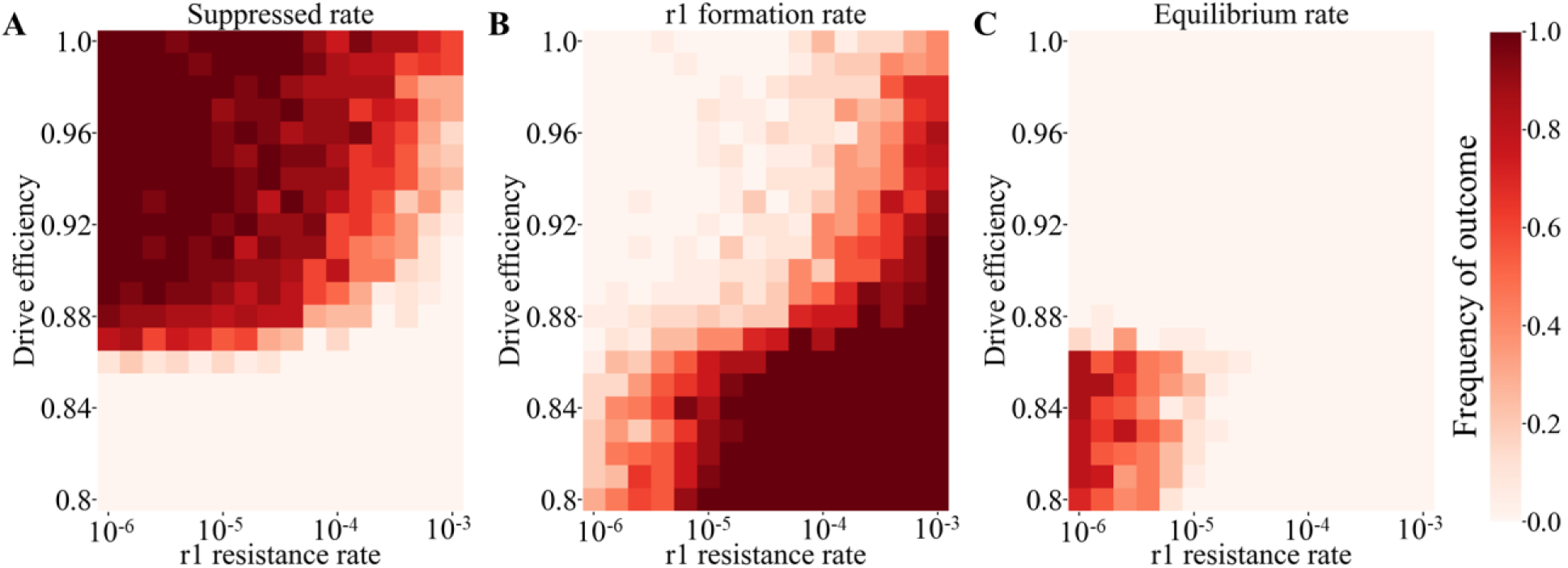
Outcomes with functional resistance in panmictic populations. Drive heterozygotes (and drive males for the haplodiploid drive) with default performance characteristics and a variable relative rate of functional resistance allele (r1) formation were released at 1% frequency into a panmictic population of 50,000 wild-type individuals. The rate of (**A**), successful suppression, (**B**) drive failure due to resistance, and (**C**) drive failure due to low genetic load were tracked. 20 simulations were assessed for each point in the parameter space. Drive loss for haplodiploids occurred in two simulations (not displayed).

**Figure S11.**
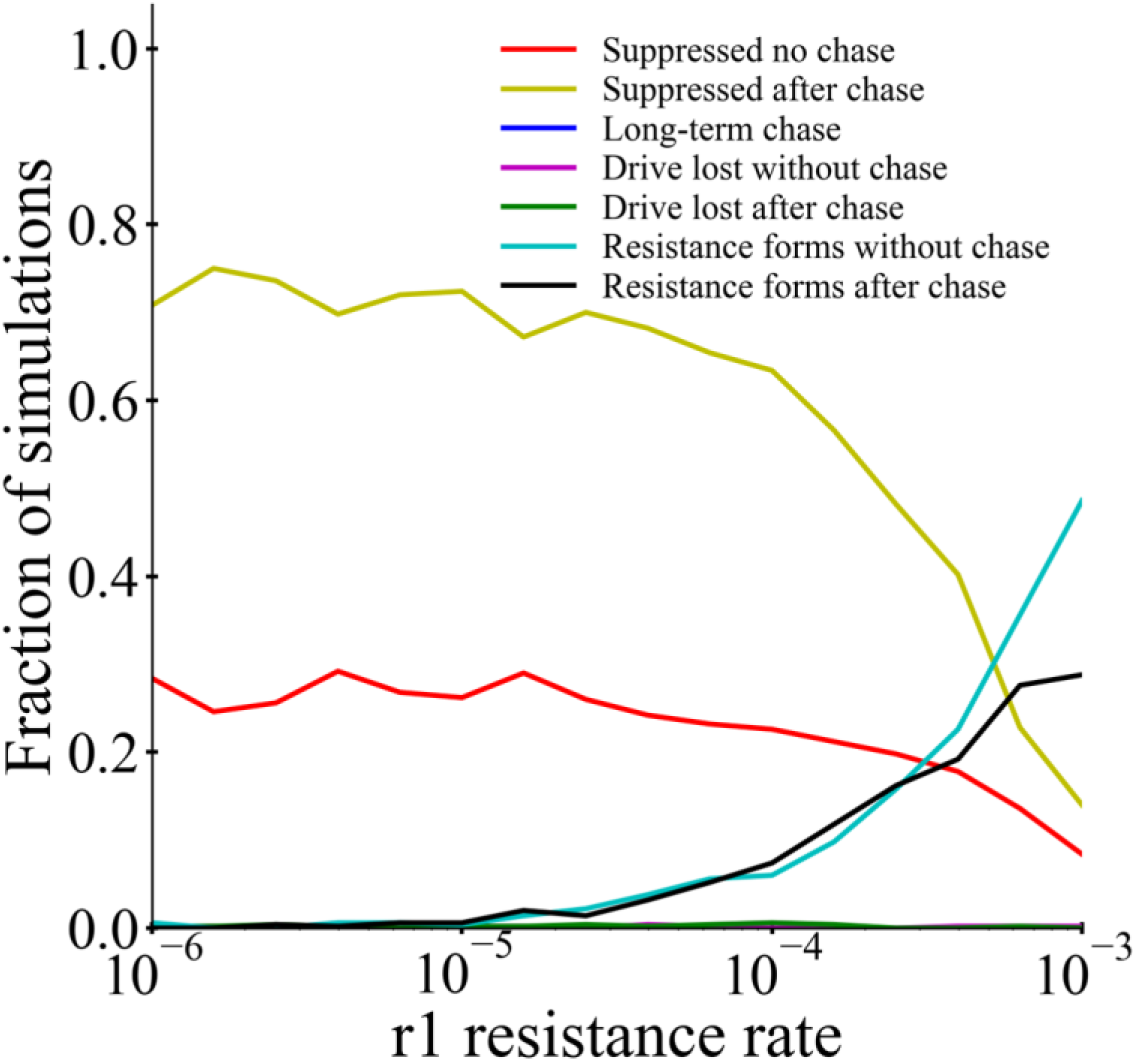
Outcomes with functional resistance in spatial populations. Female drive heterozygotes and drive males with default performance parameters and varying relative functional resistance allele (r1) formation rate were released into the middle of a spatial population of 50,000 individuals. 500 simulations were assessed for each point.

**Table S2.** Population characteristics at 90% drive frequency correlate with chasing.

|  | Haplodiploid<br>female<br>fertility<br>homing | Diploid<br>female<br>fertility<br>homing | Diploid<br>both-sex<br>fertility<br>homing | Diploid<br>driving Y<br>(X-shredder) | Diploid<br>TADS<br>Suppression |
| --- | --- | --- | --- | --- | --- |
| Population size | 3404 | 1363 | 865 | 10969 | 10999 |
| Fertile females | 10% | 7.9% | 8.1% | 9.1% | 50% |
| Sterile females | 40% | 43% | 41% | N/A | N/A |
| Fertile males | 50% | 49% | 8.8% | 8.2% | 8.3% |
| Sterile males | N/A | N/A | 42% | 83%* | 42% |
| **Expected**<br>**reproducing**<br>**females** | **345** | **114** | **12** | **103\*\*** | **934** |
| **Chasing rate** | **medium** | **medium** | **very high** | **high** | **low** |

Values are the average over 20 simulations. Except for the haplodiploid drive, data is from a previous study<sup>39</sup>. \*Drive-carrying males. \*\*Includes only females expected to mate with wild-type males.

The table shows panmictic population characteristics for the generation in which the drive is closest to 90% allele frequency in the population (using default parameters for each drive and a standard 1% release in a panmictic population of 100,000). In a spatial model, this approximately corresponds to inner edge of the drive wave of advance where population suppression is imminent. If wild-type individuals can generate offspring around here while avoiding the drive, a stochastic process, they can move into adjacent empty areas to begin a “chase”. Higher population sizes tend to reduce the frequency of this process. Thus, the both-sex fertility homing drive with only 12 reproducing females is highly prone to chasing compared to the female fertility homing drive, while the TADS suppression drive with 934 is substantially less prone to chasing. The Driving Y has a similar number of reproducing females that would be expected to generate daughters, but its total population size is overall higher, so these individuals would have fewer offspring and thus promote chasing. The haplodiploid female fertility drive, according to this analysis, would thus be somewhat better than the diploid drive at avoiding chasing, but other factors that likely affect chasing as well (particularly overall drive power/genetic load and rate of increase) result in broadly similar performance between the two drives.

## Notes

https://github.com/jchamper/ChamperLab/tree/main/Haplodiploid-Suppression-Modeling

### Competing Interest Statement

The authors have declared no competing interest.

### Summary of Updates

New data, including functional resistance alleles Fixed GitHub links Other revisions

